# Impact of ecotourism on the diversity of bacteria, archaea, and fungi in the freshwater stream of the Agua Azul Waterfalls in southeastern Mexico

**DOI:** 10.64898/2026.03.04.709505

**Authors:** Jesús Mauricio Ernesto Hernández-Méndez, Cesar Ivan Ovando-Ovando, María Emperatriz Domínguez-Espinosa, Isis Del Mazo-Monsalvo, Odín Reyes-Vallejo, Abumalé Cruz-Salomón, Michel Geovanni Santiago-Martínez

## Abstract

Freshwater streams harbor microbial communities that support ecosystem functioning. Due to their great biodiversity and geomorphological characteristics, these ecosystems are often very attractive ecotourism destinations, which makes them highly vulnerable to anthropogenic disturbances. The Agua Azul Waterfalls (Cascadas de Agua Azul, in Spanish), a major ecotourist destination located in Indigenous territories of southeastern Mexico (Chiapas, Mexico), offer a unique setting to investigate how sustained human activity influences microbial diversity and quality of water and sediments, as well as the role of local management in conserving biodiversity. To determine the sensitivity of the ecosystem to human presence, we took samples at sites encompassing gradients of ecotourism activity and performed an integrated analysis of the physicochemical properties, elemental composition, and microbial community structure of the water and sediment samples. We used more specific primers targeting the 16S rRNA genes of bacteria and archaea, and ITS2 markers for fungi. Areas associated with ecotourism activities showed changes in physicochemical parameters and microbial community composition, indicating localized human-induced impacts on this ecosystem. Although no significant differences were observed between zones, the detection of methanogenic archaea, ammonia-oxidizing archaea, sulfur-oxidizing bacteria, and saprophytic fungi in shallow sediments suggests that acetoclastic and hydrogenotrophic methanogenesis, nitrogen and sulfur cycles, cellulose degradation, and oxygen availability may influence key ecological processes within this ecosystem. Our findings highlight the potential of microbial diversity in combination with physicochemical parameters as a tool to detect early stages of human impacts on freshwater ecosystems and establish a basis for future monitoring and conservation efforts.

**Importance:** This study provides evidence of the early-stage impacts of ecotourism on biodiversity and water quality at the Agua Azul Waterfalls; a freshwater stream located in southeastern Mexico. While the water still meets basic quality standards, areas with the highest tourist activity show the first signs of nutrient enrichment (eutrophication) and changes in the potential functions performed by microbes. As the first analysis of microbial diversity in this freshwater stream, our work highlights the value of microbes as early and sensitive indicators of human impact. By directly comparing tourist and non-tourist areas, we provide evidence of how recreational pressure is transforming this ecosystem. Our findings support the great efforts of local communities to conserve this site, and we hope that these insights will guide policymakers in developing more sustainable tourism practices that preserve the biodiversity, cultural and economic value of this ecosystem, before irreversible damage occurs.

## Introduction

The Agua Azul Waterfalls (“Cascadas de Agua Azul” in Spanish) is a popular ecotourism center located in the territories of Indigenous communities (Tzeltal and Ch‘ol) in southeastern Mexico (Chiapas, Mexico). This site has become a symbol of the State of Chiapas, and is internationally known for its natural beauty, impressive waterfalls, turquoise waters and biodiversity (**Figure 1**). This site is also a very attractive tourist destination, offering a wide range of recreational activities such as swimming, diving, river rafting, boat rides, guided tours, hiking, birdwatching, among others, and welcoming approximately 300,000 visitors annually (Libert-Amico 2019). These ecotourism activities financially supports the preservation of this site and provides active income to local communities, but at the same time, visitors directly impact the ecosystem.

**Figure 1.**
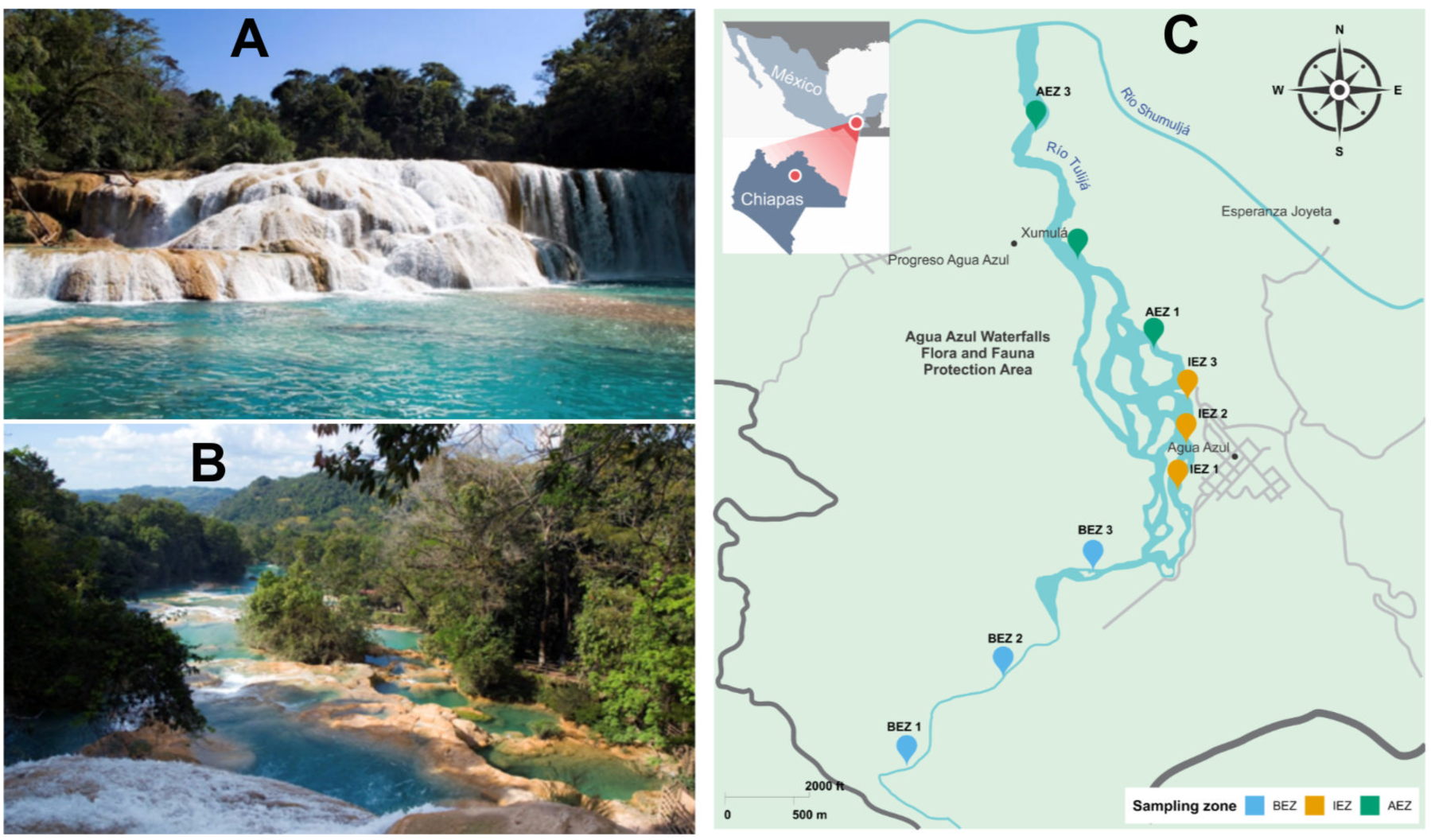
Sampling site at the Agua Azul Waterfalls (*Cascadas de Agua Azul*, in Spanish). A & B) Representative photographs taken from the CONANP website (*Comisión Nacional de Áreas Naturales Protegidas*, in Spanish; National Commission of Protected Natural Areas). C) Map of study area indicating sampling locations. Water and sediment samples were taken at three locations. Sampling zones were defined as: before ecotourism zone (BEZ), in ecotourism zone (IEZ), and after ecotourism zone (AEZ).

The Agua Azul Waterfalls (AAW) are formed by a river system that originates from three main rivers: the Otulún river, the Shumuljá river, and the Tulijá river. This river system carves a series of shallow canyons bordered by vertical limestone cliffs, creating the iconic turquoise-blue waterfalls and stepped pools characteristic of AAW. The distinctive coloration arises from the high concentrations of suspended calcium carbonate (CaCO_3_), silica (SiO_2_), magnesium hydroxide (Mg(OH)_2_), iron and aluminum oxides, combined with the naturally low levels of dissolved organic matter that allow light to scatter vividly in the water (SEMARNAT 2017).

The AAW site is classified as a flora and fauna protection area under the management of the National Commission of Natural Protected Areas (CONANP, in Spanish) (Garza-Tovar et al. 2020). This protected natural area is home to an estimated 881 species, of which 603 are vertebrate animals (18 amphibians, 20 fishes, 42 reptiles, 455 birds, and 68 mammals) and 278 are plant species. It includes 22 endemic species, representing 2.9% of the area’s biodiversity. However, 18% (159 species) of the total species are classified as at risk according to the official Mexican standard (NOM-059-SEMARNAT-2010). This record illustrates the great diversity of animals and plants; however, to date, there is no information regarding the diversity of microbes inhabiting the AAW, nor on how this microbial diversity is affected by human activities.

Microbes are essential to freshwater ecosystems, driving nutrient cycling, organic matter decomposition, and other biogeochemical processes. Because the composition and function of microbial communities respond rapidly to environmental changes, the diversity and abundance microbes can serve as sensitive indicators of ecosystem disturbances (Bai et al. 2018, Yang et al. 2020, Adedire et al. 2022). Water and sediments provide distinct but interconnected habitats, and sediments integrate environmental changes over longer periods. Human activities, including ecotourism, can alter water chemistry, nutrient inputs, sediment dynamics, and microbial communities (Almeida et al. 2007). Assessing these changes is particularly important in protected natural areas, where the economic and social benefits of tourism must be balanced with biodiversity conservation (Libert-Amico 2019).

Although there are reports on microbial diversity in natural streams in other parts of the world (URycki et al. 2020, Behera et al. 2020, Betiku et al. 2021, Qiu et al. 2020, Adedire et al. 2022), the protected natural areas of southeastern Mexico remain unexplored. Tropical freshwater streams can harbor diverse microbial communities, whose ecological responses to anthropogenic disturbances may differ from those observed in lakes and peatlands (Yang et al. 2020, Qiu et al. 2020). Furthermore, most environmental assessments focus on conventional physicochemical parameters or specific microbial groups, which limits our understanding of the ecosystem.

In this study, we investigated the diversity and abundance of bacteria, archaea, and fungi in the water and sediments of Agua Azul Falls (AAW), a tropical freshwater stream in southeastern Mexico that experiences varying levels of ecotourism activity. We integrated microbial community profiling with physicochemical and elemental characterization of the water and sediments to determine whether differences in tourism pressure are associated with changes in environmental conditions, microbial community structure, and functional potential. By comparing areas with contrasting levels of tourism activity, our study distinguishes microbial patterns associated with relatively undisturbed conditions from those associated with sustained recreational pressure.

To our knowledge, this is the first comprehensive characterization of bacterial, archaeal, and fungal diversity in both the water and sediments of AAW. Therefore, our study establishes a microbial and environmental baseline for assessing changes in this protected freshwater ecosystem. The unique characteristics of AAW make it a valuable natural environment for investigating how ecotourism shapes tropical freshwater streams and for providing a basis for conservation management strategies.

## Materials and methods

### Study sites and sample collection

The study site is the Agua Azul Waterfalls (AAW) Ecotourism Center, located in the north of the State of Chiapas, Mexico (**Figure 1**). To minimize environmental disturbance and ensure mutual benefit, the fieldwork was conducted in collaboration with representatives of the local community. The selection of sampling locations was conditioned by the complex geomorphology and the accessibility of the AAW site.

As part of this collaborative process, the research team implemented community-oriented activities aligned with local strategies for sustainable ecotourism management. These activities included workshops on solid waste management, informal discussion sessions with residents on conservation practices and environmental stewardship, and knowledge exchange focused on the ecological importance of freshwater systems. These actions were intended to support local environmental awareness while respecting community governance and Indigenous Knowledge (McAllister et al. 2025).

Water and sediment samples were collected on March 18, 2023, at three locations (three independent samples from each location) and were classified as follows: before ecotourism zone (BEZ; 17.2626 N 92.1186 W), in ecotourism zone (IEZ; 17.2525 N 92.1157 W) and after ecotourism zone (AEZ; 17.2344 N 92.1336 W). The ecotourism zone corresponds to the section of the AAW that is managed as the Cascadas de Agua Azul Ecotourism Center, where tourism-related infrastructure is in place. Therefore, this section was defined as the ecotourism zone because it represents the area with the greatest direct anthropogenic pressure associated with tourism activities. The upstream zone (BEZ) was selected as a reference location with minimal direct tourist influence, whereas the downstream zone (AEZ) was selected to evaluate the potential effects of tourism activities on water and sediment quality after the river flows through the ecotourism area. Samples were collected in early spring, prior to the onset of the rainy season and during the most intense ecotourism period. The samples from each zone were processed for analysis of physicochemical, elemental composition and microbial diversity.

Water samples (1000 mL for physicochemical analysis and 50 mL for biological analysis) were collected at each location by manually lowering a sterile bottle to a depth of approximately 20 cm following EPA-recommended procedures for monitoring recreational waters by triplicates (Betiku et al. 2021, EPA 2010). Approximately 50 g surface sediment samples were collected from the center of the stream using a stainless-steel Van Veen grab sampler (Miranda et al. 2021). The upper 3 cm of the sediment (0–3 cm depth) were subsampled from the grab under a water column of approximately 70 cm. The collected samples (50 mL of water and 50 g of sediment) were stored in previously labeled sterile bottles, placed on ice, immediately transferred to the laboratory and kept frozen at-20 °C until subsequent biological analysis.

### Analytical methods

Physicochemical parameters of water samples were analyzed according to Standard Methods for Examination of Water and Wastewater (APHA 2012) using 1000 mL of sample. The parameters analyzed were color, turbidity, electrical conductivity (EC), total solids (TS), total suspended solids (TSS), total volatile solids (TVS), total dissolved solids (TDS), pH, alkalinity, acidity, chemical oxygen demand (COD), total Kjeldahl nitrogen (TKN), and total phosphorus (TP). Also, the concentrations of metals (Ca, Mg, Na, K) were analyzed using the standard EPA method (6010) with an inductively coupled plasma optical emission spectrophotometer (ICP-OES, Optima 7000 DV, PerkinElmer, United States).

Triplicate sediment samples from the same sampling zones were analyzed for elemental composition and morphological structure. Sediment samples were air-dried, sieved, and ground to a particle size below 2 mm using a Rocklabs grinder for one minute. The processed samples were characterized using a Hitachi S-5500 field-emission scanning electron microscope (FE-SEM; Hitachi High-Technologies, Japan) coupled with energy-dispersive X-ray spectroscopy (EDS; Oxford Instruments, UK) operating at an accelerating voltage of 5 kV for high-resolution imaging (Ramírez-Galdámez et al. 2024). SEM micrographs were obtained at low (≈250×) and high (≈10,000×) magnifications, and EDS spectra were collected from representative areas to determine the relative abundance of major and minor elements.

All physicochemical and elemental parameters of the samples were measured during the following 24 hours in triplicate. The data were statistically analyzed using a one-way ANOVA followed by Tukey test to determine if there were significant differences between the different study zones (BEZ, IEZ and AEZ). The values were considered significantly different at *p* < 0.05 (*), *p* < 0.01 (**), *p* < 0.001(***) and *p* < 0.0001 (****). Before the analysis, the assumptions of the ANOVA were thoroughly verified.

Within-zone physicochemical variability was additionally assessed using multivariate analyses of the measured water properties. Variables were centered and standardized to unit variance prior to analysis. Principal component analysis (PCA) was performed on the standardized data to visualize multivariate physicochemical patterns among replicate samples and sampling zones. Within-zone variability was quantified as the Euclidean distance of each sample to the centroid of its respective sampling zone in standardized multivariate space. Homogeneity of multivariate dispersion among BEZ, IEZ, and AEZ was evaluated using betadisper, followed by a permutation test with 9,999 permutations. When the global test was significant, pairwise differences in dispersion were evaluated using Tukey-adjusted post hoc comparisons. Analyses were conducted in R (v4.6.1) using the vegan package (v2.7-5). (*See* Data and code availability statement).

### DNA extraction, Library construction and sequencing

Water samples and a subsample of the collected sediment was used for total DNA extraction according to the manufacturer’s protocol. The water sample (50 mL) was filtered using a Cobetter® capsule filter (Cobetter Filtration Equipment Co. Ltd, China) with sterile polyethersulfone (PES) filter membranes measuring 47 mm in diameter and having a pore size of 0.22 µm. The sediment sample was agitated for 5 minutes using a vortex (GENIE 2, DG-3030A2) at medium speed, and a 250 mg subsample was taken from each homogenized sediment sample for DNA extraction. Subsequently, the ZymoBiomics^TM^ DNA Miniprep commercial kit (Zymo Research Corporation, California, USA) was used for the extraction of total DNA from both the water and sediment samples following the manufacturer’s protocol. Total DNA was quantified using a Qubit dsDNA BR (Broad Range) Assay Kit (Invitrogen).

The extracted total DNA was used as a template for PCR amplification. Bacterial 16S rRNA gene fragments (V4–V5 region) were amplified using primers 515F (5′-GTG YCA GCM GCC GCG GTA A-3′) and 926R (5′-CC GYC AAT TYM TTT RAG TTT-3′). Archaeal 16S rRNA gene fragments (V6–V8 region) were amplified using primers 956F (5′-TYA ATY GGA NTC AAC RCC-3′) and 1401R (5′-CR GTG WGT RCA AGG RGC A-3′) (Peña-Ocaña et al. 2022). Fungal communities were characterized by amplifying the ITS2 region using primers fITS7 (5′-GTG ART CAT CGA ATC TTT G-3′) and ITS4 (5′-TCC TCC GCT TAT TGA TAT GC-3′) (Ihrmark et al. 2012, Kozich et al. 2013). Barcoded DNA libraries were prepared and analyzed at the UConn Center for Genome Innovation (CGI) facility (Storrs, Connecticut, USA) with Illumina paired-end library preparation and 250 bp paired-end sequencing on an Illumina MiSeq (Illumina, Inc. CA, USA).

### Microbial marker-gene amplicon data analysis

#### Microbial marker-gene sequence processing

Raw paired-end amplicon sequences were processed using the nf-core/ampliseq pipeline (v2.15.0) implemented in Nextflow (v25.10.3). For bacterial and archaeal 16S rRNA gene datasets, primer sequences were removed using Cutadapt (v4.6) with double-primer detection enabled, a minimum overlap of 10 bp, and a maximum error rate of 0.12. Amplicon sequence variants (ASVs) were inferred using DADA2 (v1.30.0), including quality filtering, error correction, dereplication, paired-end merging, and chimera removal. Bacterial reads were truncated to 230 bp (forward) and 220 bp (reverse), whereas archaeal reads were truncated to 230 bp for both forward and reverse reads. Denoising was performed using maxEE = 2 for bacterial datasets and maxEE = 3 for archaeal datasets, with truncQ = 2. Archaeal read-tracking statistics indicated that 80.4–91.8% of denoised forward reads were successfully merged, confirming adequate paired-end overlap.

For fungal ITS datasets the reads were quality-filtered using fastp (v0.24.0) with a sliding-window approach (window size = 4 bp; minimum mean Phred score=20) (Kyle and Klassen 2025), trimming both read ends and enabling paired-end adapter detection. Filtered reads were subsequently processed with nf-core/ampliseq, where ITS2 sequences were extracted (cut_its = ITS2) and denoised with DADA2 without additional read truncation (trunclenF=0, trunclenR=0, maxEE=2, truncQ=2).

#### Taxonomic assignment

Taxonomic assignment of bacterial and archaeal ASVs was performed using the SILVA reference database (v138.2), whereas fungal ITS2 ASVs were classified against the UNITE reference database (v10.0) (Abarenkov et al. 2024). Resulting ASV tables were imported into phyloseq (v1.56.0) for downstream analyses. Prior to diversity analyses, low-abundance ASVs were removed by excluding taxa represented by fewer than 10 total reads across the dataset or detected in fewer than two samples.

#### Diversity analysis

Downstream statistical analyses were performed in R (v4.6.1). Alpha diversity was calculated from rarefied ASV tables generated using rarefy_even_depth() in phyloseq (McMurdie and Holmes, 2013), with samples randomly subsampled without replacement to the minimum sequencing depth within each dataset. Observed richness, Chao1 richness, Shannon diversity, and Simpson diversity were calculated using estimate_richness(). Differences in alpha-diversity metrics among sampling zones were evaluated using Kruskal–Wallis tests.

For beta-diversity analyses, ASV counts were transformed to relative abundances using transform_sample_counts(), and Bray–Curtis dissimilarities were calculated using distance() in phyloseq. Principal coordinates analysis (PCoA) was used to ordinate Bray–Curtis dissimilarities, and ordinations were visualized using ggplot2 (Wickham 2016). Differences in community composition among sampling zones were evaluated by permutational multivariate analysis of variance (PERMANOVA) using adonis2() in vegan (v2.7-5), with 999 permutations. Homogeneity of multivariate dispersion among sampling zones was assessed using betadisper(), with significance evaluated by a permutation test using 999 permutations.

#### Differential abundance analyses

Differential abundance was primarily assessed using DESeq2 (v1.52.0) at the genus level (Anderson et al. 2006). Analyses were performed on filtered, non-rarefied count data using negative binomial generalized linear models with the design formula ∼ zone. Size factors were estimated using sfType = “poscounts”. Pairwise contrasts were performed for BEZ versus AEZ, IEZ versus AEZ, and IEZ versus BEZ. P-values were adjusted for multiple testing using the Holm procedure, and genus-level features with an adjusted P < 0.05 were considered differentially abundant.

As a complementary approach, differential abundance was evaluated using ANCOM-BC2 (v2.14.0) at the genus level on filtered, non-rarefied count data. Analyses included global and pairwise tests among sampling zones, structural-zero detection, and pseudo-count sensitivity analysis (pseudo_sens = TRUE). P-values were adjusted using the Holm procedure, with adjusted P < 0.05 considered statistically significant. Taxa detected in the primary analysis were distinguished from those remaining robust after pseudo-count sensitivity analysis.

#### Environmental association analyses

Relationships between microbial community composition and measured environmental variables were evaluated using distance-based redundancy analysis (dbRDA) implemented with the dbrda() function in the vegan package. Community dissimilarities were calculated using Bray–Curtis distances, and environmental variables were aligned with the corresponding biological samples before analysis. The significance of the constrained ordination model and individual environmental variables was assessed using permutation tests (999 permutations).

#### Functional and ecological prediction analyses

Functional profiles were predicted from bacterial (water and sediment) and archaeal (sediment) 16S rRNA gene ASVs using PICRUSt2 (v2.5.3) (Douglas et al. 2019). Predicted MetaCyc metabolic pathways were normalized to relative abundances within each sample by dividing the predicted abundance of each pathway by the total predicted pathway abundance. Differences in pathway abundances among sampling zones were evaluated independently for each MetaCyc pathway using the Kruskal–Wallis test. P-values were adjusted for multiple comparisons using the Benjamini–Hochberg false discovery rate (FDR), and adjusted P values < 0.05 were considered statistically significant. To infer the ecological roles of fungal taxa, functional guild annotation was performed using FUNGuild (Nguyen et al. 2016). Functional annotation was executed using the default fungal database and the output included predicted trophic modes, functional guilds, and the confidence ranking. Only assignments with confidence levels of Probable or Highly Probable were considered for downstream ecological interpretation.

## Results

### Physicochemical characteristics of water

The results of the physicochemical analysis of surface water samples (**Figure 2**) show a significant difference among sampling zones for several measured variables. One-way ANOVA detected significant differences in alkalinity, calcium (Ca), color, electrical conductivity, pH, potassium (K), total dissolved solids (TDS), total Kjeldahl nitrogen (TKN), total solids (TS), turbidity and total volatile solids (TVS) (all *p* < 0.05), with relatively large effect sizes (η² = 0.71–0.99). Tukey comparisons showed that IEZ had higher Ca, color, electrical conductivity, pH, TDS, TKN, TS, turbidity and TVS than BEZ, whereas alkalinity was lower in IEZ than in BEZ. Several variables also differed between IEZ and AEZ, including Ca, color, pH, TNK and turbidity, while BEZ and AEZ differed in alkalinity, color, pH, K, and TS. No significant differences were detected for COD, Mg, Na, TP, and TSS (*p* > 0.05), although COD and TSS showed comparatively large, estimated effect sizes (η² = 0.61 and 0.56, respectively). These results motivated us to evaluate the diversity, abundance, and potential metabolic functions of the associated microbes, since these factors could affect microbial metabolism, particularly those that depend on the availability of light and oxygen.

**Figure 2.**
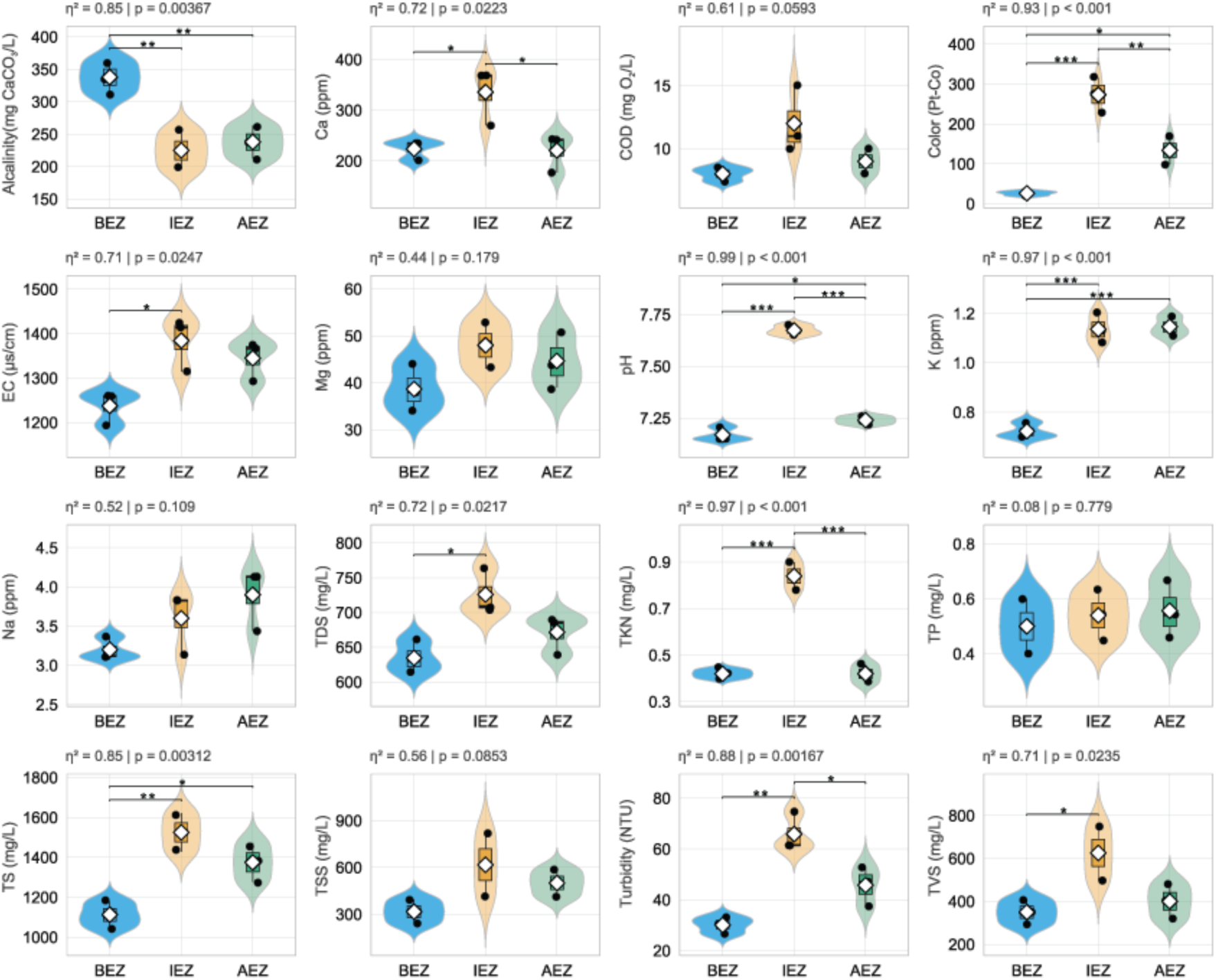
Physicochemical characterization of water samples from the Agua Azul Waterfalls across sampling zones. Sampling zones were defined as: before the ecotourism zone (BEZ), within the ecotourism zone (IEZ), and after the ecotourism zone (AEZ). Parameters evaluated included alkalinity, calcium (Ca), chemical oxygen demand (COD), color, electrical conductivity (EC), magnesium (Mg), pH, potassium (K), sodium (Na), total dissolved solids (TDS), total Kjeldahl nitrogen (TKN) and total phosphorus (TP). Violin plots show the distribution of observations for each sampling zone, with superimposed boxplots representing the median and interquartile range (IQR). Individual observations are shown as jittered points, and white diamonds indicate the mean (n = 3 per sampling zone). Differences among sampling zones were evaluated using one-way ANOVA followed by Tukey’s HSD post hoc test for pairwise comparisons. The overall *p* value and effect size (η²) from the one-way ANOVA are displayed above each panel. Significant pairwise differences determined by Tukey’s HSD test are indicated by horizontal brackets and asterisks: *p* < 0.05 (*), *p* < 0.01 (**), and *p* < 0.001 (***).

Principal component analysis (PCA) of standardized water physicochemical variables further showed zone-associated multivariate structuring among replicate samples. PC1 and PC2 accounted for 62.7% and 12.9% of the total variance, respectively (75.6% cumulatively). Replicates from BEZ, AEZ, and IEZ occupied non-overlapping ranges along PC1, with scores ranging from-4.25 to-2.93,-0.53 to 0.17, and 3.64 to 3.75, respectively. The largest absolute loadings on PC1 corresponded to color (0.305), turbidity (0.294), pH (0.293), total solids (0.286), and TKN (0.274). Multivariate dispersion differed among sampling zones (permutation test, *F*_2,6_ = 12.38, *p* = 0.0073), with mean distances to zone centroids of 1.49 ± 0.27, 1.96 ± 0.25, and 2.42 ± 0.15 for BEZ, AEZ, and IEZ, respectively. Post hoc comparisons showed greater dispersion in IEZ than in BEZ (Tukey-adjusted *P* = 0.006), whereas AEZ did not differ significantly from either zone (adjusted *p* > 0.10; **Figure S1**).

### Sediment elemental composition

The elemental composition of sediments showed a consistent predominance of oxygen (45– 55%), carbon (15–20%), silicon (12–20%), calcium (15–22%), and magnesium (4–8%) across the three sampling zones (BEZ, IEZ, AEZ) (**Figure 3**).

**Figure 3.**
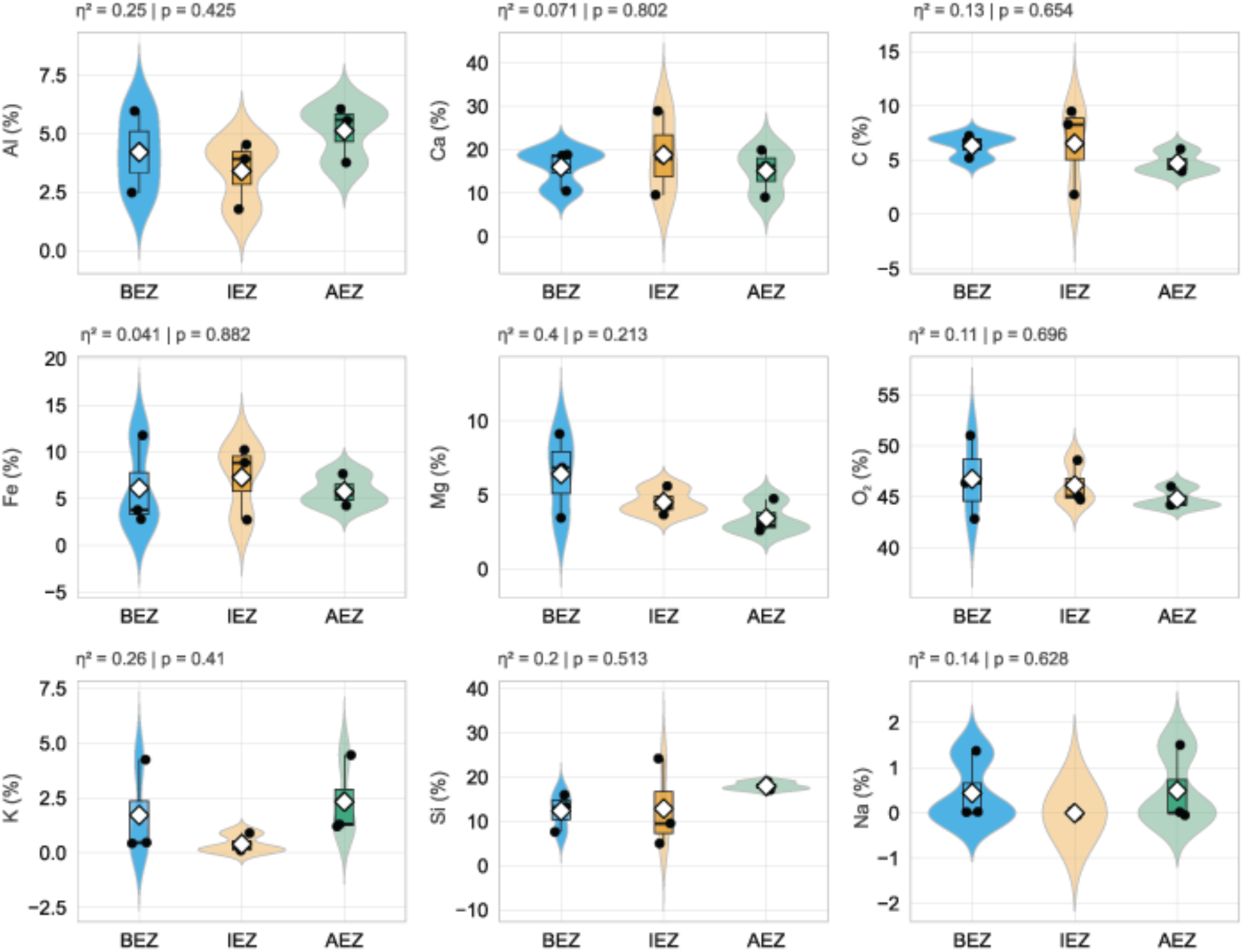
The elemental characterization of sediment samples from the Agua Azul Waterfalls. Sampling zones were defined as: before ecotourism zone (BEZ), in ecotourism zone (IEZ), and after ecotourism zone (AEZ). Parameters evaluated included aluminum (Al), calcium (Ca), carbon (C), iron (Fe), magnesium (Mg), oxygen (O_2_) potassium (K), silicon (Si) and sodium (Na). Violin plots show the distribution of observations for each sampling zone, with superimposed boxplots representing the median and interquartile range (IQR). Individual observations are shown as jittered points, and white diamonds indicate the mean (n = 3 per sampling zone). Differences among sampling zones were evaluated using one-way ANOVA followed by Tukey’s HSD post hoc test for pairwise comparisons. The overall p value and effect size (η²) from the one-way ANOVA are displayed above each panel.

In contrast to the water physicochemical characteristics, sediment elemental composition did not differ significantly among BEZ, IEZ, and AEZ. One-way ANOVA detected no significant differences for any of the nine measured elements (all *p* > 0.05), including oxygen, calcium, carbon, magnesium, silicon, iron, aluminum, potassium, and sodium. Accordingly, no significant pairwise differences among sampling zones were identified by the Tukey post hoc comparisons. Estimated effect sizes varied among elements, with magnesium exhibiting the largest value (η² = 0.40), followed by potassium (η² = 0.26), aluminum (η² = 0.25), and silicon (η² = 0.20), although none of these differences reached statistical significance. The remaining elements showed smaller estimated effect sizes (η² = 0.04–0.14). Overall, the distributions displayed substantial within-zone variability for several elements, particularly magnesium, potassium, and silicon, without statistically supported differences in elemental composition among the three sampling zones.

The field-emission scanning electron microscopy (FE-SEM) micrographs showed clear morphological differences among sediment samples from the three zones (**Figure S2**). In BEZ samples, particles exhibited compact surfaces, smooth crystalline edges, and well-defined crystalline structures. IEZ samples displayed highly fragmented particles with irregular, angular edges, increased surface roughness, and heterogeneous grain size distributions. The AEZ samples exhibited irregular agglomerated aggregates.

### Bacterial community composition and diversity across water and sediment

The bacterial water dataset comprised nine samples (three each from BEZ, IEZ, and AEZ), with 431,669 reads and 7,670 ASVs before filtering. After filtering, all samples were retained, comprising 262,778 reads and 1,114 ASVs. No significant differences in alpha diversity were detected among zones for observed richness, Chao1 richness, Shannon diversity, or Simpson diversity (Kruskal–Wallis, *H* = 5.956, df = 2, *P* = 0.0509 for all metrics) **(Figure S3-B)**. No significant differences in Bray–Curtis community composition were detected among BEZ, IEZ, and AEZ (PERMANOVA, *F*_2,6_ = 2.138, R² = 0.416, *P* = 0.066), and multivariate dispersion did not differ significantly among zones (betadisper, *F*_2,6_ = 1.465, *p* = 0.336) **(Figure S3-C)**.

At the genus level, DESeq2 identified 151 differentially abundant features between BEZ and AEZ, 73 between IEZ and AEZ, and only one between IEZ and BEZ. Differential abundance was therefore strongly concentrated in comparisons involving AEZ, whereas IEZ and BEZ showed comparatively little differentiation. Several genera, including *Dinghuibacter*, *Thermomonas*, and *Sphingorhabdus*, were significantly different in both BEZ–AEZ and IEZ–AEZ comparisons, while *Sphingobium* was the only genus significantly differentiating IEZ from BEZ (**Table S2)**. In the complementary ANCOM-BC2 analysis, two primary hits were detected globally and two for BEZ versus AEZ, whereas no differentially abundant taxa were detected for the other pairwise contrasts.

The 16S rRNA gene bacterial sediment dataset comprised nine samples and 1,295 taxa after filtering. Alpha diversity did not differ significantly among BEZ, IEZ and AEZ for observed richness, Chao1 richness (Kruskal–Wallis, *H* = 2.40, *df* = 2, *P* = 0.301 for both metrics), Shannon diversity or Simpson diversity (*H* = 3.82, *df* = 2, *P* = 0.148 for both metrics) (**Figure S4-B)**. Bacterial community composition differed significantly among BEZ, IEZ and AEZ based on Bray–Curtis dissimilarities (PERMANOVA, *F*_2,6_ = 1.89, R² = 0.387, *p* = 0.018). This result was not accompanied by significant differences in multivariate dispersion (betadisper, *F*_2,6_ = 0.125, *p* = 0.871) (**Figure S4-C)**.

At the genus level, 33 unique differentially abundant features were identified by DESeq2 across pairwise comparisons after Holm correction, with 14 detected between BEZ and AEZ, 23 between IEZ and AEZ, and five between IEZ and BEZ. Most differential abundance was associated with comparisons involving AEZ, whereas relatively few features differentiated IEZ and BEZ (**Table S3)**. Several genus-resolved features, including *Thiobacillus*, *Flavihumibacter*, *Terrimonas*, *Sphingomonas*, and *Lacibacter*, were consistently detected in both BEZ–AEZ and IEZ–AEZ comparisons. In contrast, ANCOM-BC2 detected a single genus in the global test, which remained robust after pseudo-count sensitivity analysis, whereas no differentially abundant taxa were detected in any pairwise comparison (BEZ vs AEZ, IEZ vs AEZ, or IEZ vs BEZ).

When water and sediment samples were considered together, the relative abundance profiles of the dominant bacterial taxa showed heterogeneous patterns across matrices and sampling zones (**Figure 4-A**). A substantial fraction of the bacterial community remained unclassified in both water and sediment, with mean relative abundances of 29.0% and 27.6%, respectively. Among the classified genera, *Sandaracinomonas* showed the strongest descriptive difference between matrices, with a higher mean relative abundance in water (15.3%) than in sediment (5.2%), whereas *Flavobacterium* showed the opposite tendency (4.8% in water and 7.9% in sediment). Considerable variation was nevertheless observed among samples and sampling zones. Accordingly, none of the taxa for which an overall water–sediment comparison was available showed a significant difference between matrices after Benjamini-Hochberg correction (Wilcoxon tests, all *q* = 0.895). *Acinetobacter*, *Exiguobacterium*, and *Thermomonas* were represented only in sediment, whereas *Brevundimonas* was represented only in water, precluding corresponding matrix-level comparisons.

**Figure 4.**
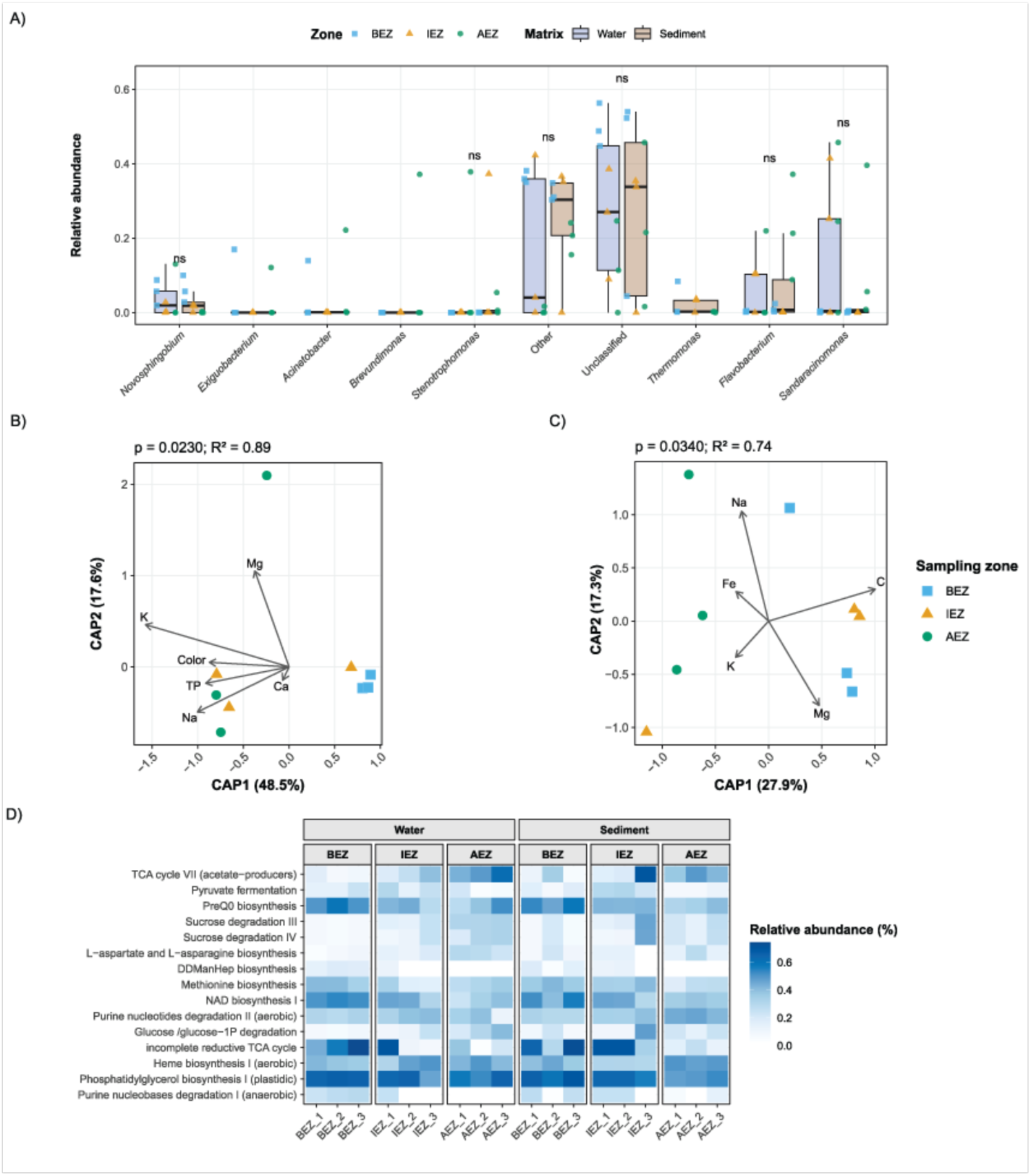
Taxonomic composition, environmental associations, and predicted functional profiles of bacterial communities in water and sediment. **A)** Top genus with highest mean relative abundance across water and sediment samples. Boxplots shows relative abundance distributions by sample matrix, with individual observations overlaid as jittered points in each zone (BEZ, IEZ, and AEZ). Differences between water and sediment were evaluated separately for each taxon using Wilcoxon rank-sum tests, with p-values adjusted for multiple comparisons using the Benjamini– Hochberg (BH) procedure. ns indicates q ≥ 0.05. (B–C) Distance-based redundancy analysis (dbRDA) of bacterial community composition in water (B) and sediment (C), based on Bray–Curtis dissimilarities and constrained by measured environmental variables. Points represent individual samples and colors indicate sampling zones. Arrows show significant environmental variables (p ≤ 0.05; up to eight variables per panel). Overall model p-value and R² are displayed above each ordination. D) Predicted bacterial MetaCyc pathway profiles across water and sediment. Top pathways shared between matrices. Differences between sampling zones were evaluated separately within water and sediment. Sampling zones: before the ecotourism zone (BEZ), within the ecotourism zone (IEZ), and after the ecotourism zone (AEZ).

### Environmental drivers of bacterial community composition

Given the spatial variation observed in bacterial community composition, particularly in sediment and in taxonomic comparisons involving AEZ, we next evaluated the extent to which environmental variables were associated with community structure using distance-based redundancy analysis (dbRDA). Water and sediment communities were analyzed separately to account for matrix-specific environmental conditions (**Figure 4-B, C**).

In water samples, dbRDA revealed a significant association between the measured environmental variables and bacterial community composition (permutation test, *p* = 0.023). Although no individual environmental variable reached statistical significance, potassium, magnesium, color, and total phosphorus showed the strongest associations with community structure (*p* = 0.056-0.070). The ordination further showed variation in the distribution of samples from the three sampling zones along the environmental gradients, with BEZ samples clustering toward positive CAP1 values and greater dispersion observed among AEZ samples.

In sediment samples, dbRDA similarly revealed a significant association between the measured environmental variables and bacterial community composition (permutation test, *p* = 0.034; Fig. 3C). In contrast to water, carbon and magnesium were significantly associated with sediment bacterial community structure (*p* = 0.004 and *p* = 0.024, respectively), while sodium showed a weaker, non-significant association (*p* = 0.062). The ordination showed that most BEZ and IEZ samples were positioned toward positive CAP1 values, in the direction associated with increasing carbon, whereas AEZ samples were predominantly distributed toward negative CAP1 values.

### Predicted bacterial functional profiles across sampling zones

Given the taxonomic and environmentally associated variation observed across bacterial communities, we examined whether these patterns were accompanied by differences in their predicted functional potential. PICRUSt2-based functional predictions showed comparable weighted NSTI values for sediment (mean ± SD: 0.151 ± 0.102, n = 9) and water (0.175 ± 0.104, n = 9), providing context for the phylogenetic proximity underlying the functional predictions. Functional profiling identified a set of 15 MetaCyc pathways prioritized across bacterial communities from water and sediment **(Figure 4-D**). These pathways encompassed functions related to central carbon metabolism, fermentation, amino acid and nucleotide metabolism, and cofactor and membrane-component biosynthesis. Among the highest-ranked pathways were TCA cycle VII (acetate producers), pyruvate fermentation to acetate and lactate II, preQ₀ biosynthesis, and sucrose degradation pathways. Despite variation in their predicted relative abundances among sampling zones, none of the 15 prioritized pathways differed significantly among BEZ, IEZ, and AEZ after Benjamini–Hochberg correction, either in water or sediment (all FDR-adjusted *p* > 0.05).

### Archaeal community composition and taxonomic patterns in sediment

Water samples yielded no archaeal or ITS fungal features after quality filtering and taxonomic processing and were therefore excluded from downstream analyses of these microbial groups. The 16S rRNA gene archaeal sediment dataset comprised nine samples and 658 taxa after filtering. No significant differences were detected among BEZ, IEZ and AEZ for observed richness (Kruskal–Wallis, *H* = 4.73, *df* = 2, *p* = 0.094), Chao1 richness (*H* = 5.07, *df* = 2, *P* = 0.079), Shannon diversity or Simpson diversity (*H* = 5.42, *df* = 2, *P* = 0.066 for both metrics) **(Figure S5-B**). Archaeal community composition differed significantly among BEZ, IEZ and AEZ based on Bray–Curtis dissimilarities (PERMANOVA, *F*_2,6_ = 6.09, R² = 0.670, *p* = 0.005) **(Figure S5-C**), while no significant differences were detected in multivariate dispersion (betadisper, *F*_2,6_ = 2.10, *p* = 0.214).

At the genus level, DESeq2 identified 18 unique differentially abundant features across pairwise comparisons, with 15 detected between BEZ and AEZ, 12 between IEZ and AEZ, and only two between IEZ and BEZ **(Table S4**). Several genus-resolved features, including *Methanothrix*, *Methanospirillum*, *Methanoregula*, *Nitrosarchaeum*, and *Candidatus Nitrosotenuis*, were differentially abundant in both BEZ–AEZ and IEZ–AEZ comparisons. ANCOM-BC2 detected 10 primary hits in the global test, four of which remained robust after pseudo-count sensitivity analysis. Pairwise testing yielded seven primary hits for BEZ versus AEZ, none of which remained robust, and 11 for IEZ versus AEZ, of which one remained robust.

Consistent with this pronounced spatial heterogeneity, the relative abundance profiles of the dominant archaeal taxa showed distinct distributions among sampling zones **(Figure 5-A**). *Nitrosarchaeum* (ammonia-oxidizing archaea) showed higher relative abundances in IEZ, whereas *Methanothrix* (methane-producing anaerobic archaea) was consistently more abundant in AEZ. Other ammonia-oxidizing archaea, including *Candidatus Nitrosocosmicus*, *Candidatus Nitrososphaera*, and *Candidatus Nitrosotenuis*, also displayed variable distributions among zones, while a substantial fraction of the archaeal community remained taxonomically unclassified. When the selected dominant taxa were evaluated individually using Kruskal–Wallis tests, however, none remained significant after Benjamini–Hochberg correction (all *q* ≥ 0.083).

**Figure 5.**
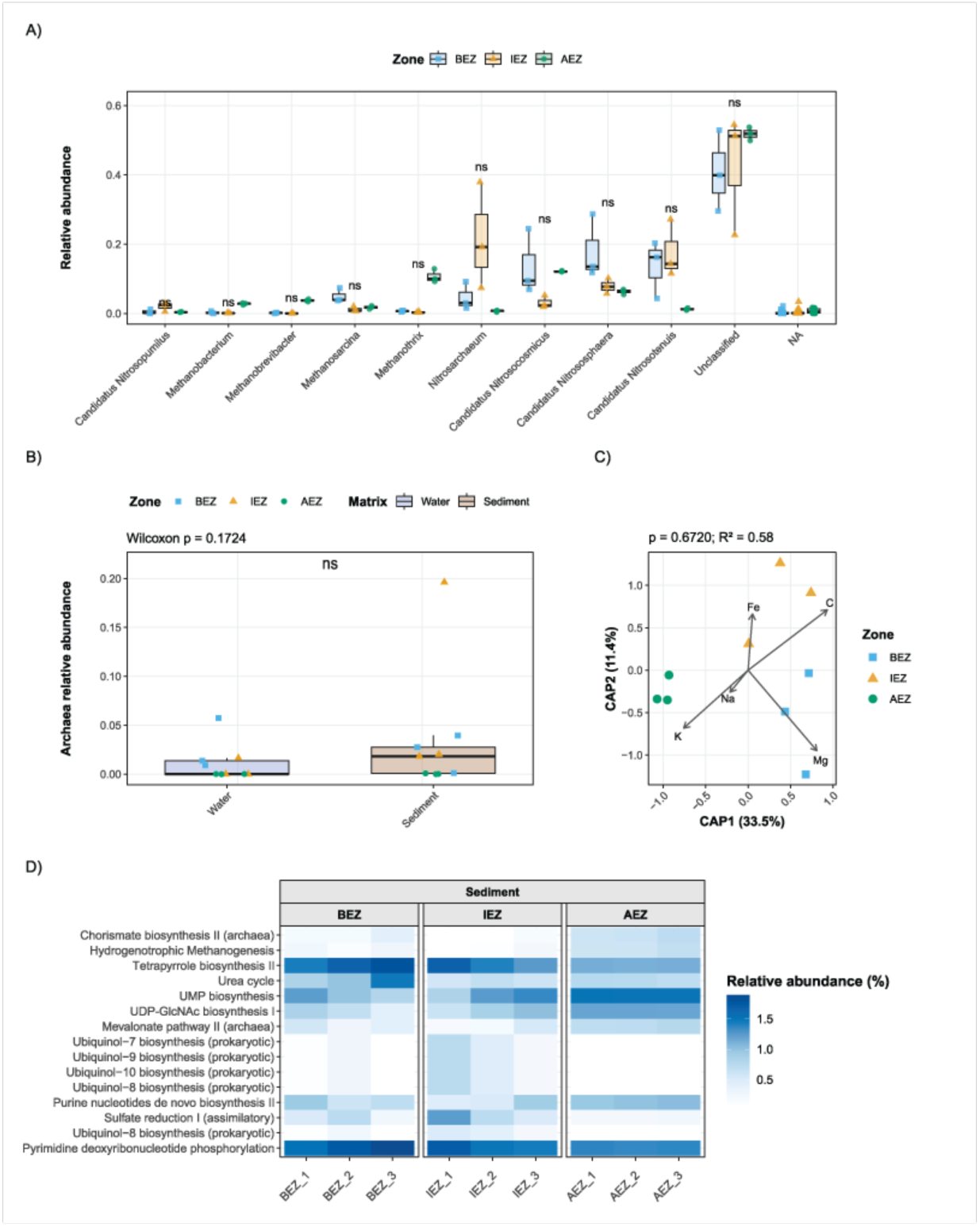
Taxonomic composition, environmental associations, and predicted functional profiles of archaeal communities in sediment. **A**) Boxplots of top genus with the highest mean relative abundance across zone (BEZ, IEZ, and AEZ) with individual observations overlaid as jittered points. Differences between zone were evaluated separately for each taxon using Wilcoxon rank-sum tests, with p-values adjusted for multiple comparisons using the Benjamini–Hochberg (BH) procedure. ns indicates q ≥ 0.05. **B)** Archaeal relative abundance in bacterial 16S datasets across water and sediment. Boxplots show the distribution of archaeal relative abundance in each sample matrix, with individual samples overlaid as jittered points in each zone (BEZ, IEZ, and AEZ). Differences between water and sediment were evaluated using a Wilcoxon rank-sum test. ns indicates no significant difference (p ≥ 0.05). **C**) Distance-based redundancy analysis (dbRDA) of archaeal community composition in sediment, based on Bray–Curtis dissimilarities and constrained by measured environmental variables. Points represent individual samples. Arrows show significant environmental variables (p ≤ 0.05; up to eight variables per panel), and the overall model p-value and R² are displayed above each ordination. **D**) Predicted bacterial MetaCyc pathway profiles across water and sediment. Differences among sampling zones were evaluated separately within water and sediment. Asterisks next to pathway names indicate pathways significant in at least one sample matrix after FDR correction. Sampling zones: before the ecotourism zone (BEZ), within the ecotourism zone (IEZ), and after the ecotourism zone (AEZ).

As a complementary analysis, sequences assigned to archaea within the bacterial 16S rRNA gene datasets were examined separately to provide an exploratory comparison of the archaeal signal between water and sediment **(Figure 5-B**). This approach allowed both matrices to be evaluated using archaeal-assigned sequences recovered under the same bacterial 16S rRNA gene processing framework, independently of the sediment-specific archaeal dataset described above. Archaeal-assigned sequences represented a generally small and highly variable fraction of the bacterial 16S rRNA gene profiles, with several samples containing no detectable archaeal signal. Although higher proportions occurred in some sediment samples, particularly within IEZ, the overall archaeal proportion did not differ significantly between water and sediment (Wilcoxon rank-sum test, *p* = 0.172). Thus, the bacterial 16S rRNA gene datasets provided no statistical evidence of a matrix-associated difference in the recovered archaeal signal.

### Environmental associations with archaeal community composition

To evaluate whether the spatial variation observed in sediment archaeal communities was associated with the measured environmental conditions, we performed distance-based redundancy analysis (dbRDA; **Figure 5-C**). The overall environmental model was not statistically significant, and none of the individual sediment variables showed a significant association with archaeal community composition (permutation tests, all *p* ≥ 0.303). Although the ordination displayed spatial structuring of the samples, with AEZ positioned primarily toward negative CAP1 values and IEZ and BEZ predominantly toward positive values, this pattern was not statistically supported by the measured environmental predictors. Thus, the environmental variables included in the analysis did not provide significant explanatory support for the zone-associated differences in archaeal community composition detected by PERMANOVA.

### Predicted functional profiles of archaeal communities in sediment

Functional predictions of sediment archaeal communities showed weighted NSTI values comparable to bacterial communities (mean ± SD: 0.176 ± 0.016, n = 9). Functional profiling identified 15 prioritized MetaCyc pathways showing variable predicted abundances across sampling zones **(Figure 5-D**). These pathways encompassed functions related to central biosynthetic processes, methanogenesis, nitrogen metabolism, nucleotide and cofactor biosynthesis, and assimilatory sulfate reduction. Among the highest-ranked pathways, archaeal chorismate biosynthesis and hydrogen-dependent CO_2_-reducing methanogenesis (hydrogenotrophic methanogenesis) showed higher median predicted abundances in AEZ, whereas the urea cycle showed its highest median abundance in BEZ. Several ubiquinol biosynthesis pathways displayed higher predicted abundances in IEZ. Although some of these pathways showed relatively large effect sizes (0.66–0.87) and nominal Kruskal–Wallis *p* values below 0.05, none remained significant after Benjamini–Hochberg correction (all adjusted *p* = 0.257 among the 15 prioritized pathways). Thus, no statistically supported zone-associated differences were detected among the prioritized predicted archaeal functions.

### Fungal community diversity and composition

The ITS sediment dataset comprised six samples distributed across BEZ, IEZ and AEZ, with 314 taxa retained after filtering. Alpha diversity did not differ significantly among zones for observed richness, Chao1 richness, Shannon diversity or Simpson diversity (Kruskal–Wallis, *H* = 4.57, *df* = 2, *p* = 0.102 for all metrics) **(Figure S6-B**).

Fungal community composition based on Bray–Curtis dissimilarities did not differ significantly among zones (PERMANOVA*, F*_2,6_ = 2.71, R² = 0.643, *p* = 0.0667) (**Figure S6-C)**. However, multivariate dispersion differed significantly among zones (betadisper, *F*_2,6_ = 7.67 × 10²⁹, *p* = 0.00139), indicating substantial heterogeneity in within-zone dispersion and warranting caution in interpreting the PERMANOVA result.

At genus level, DESeq2 identified one unique differentially abundant feature. *Acrocalymma*was more abundant in BEZ than AEZ (log2 fold change = 23.42, *Wald z* = 5.91, *adjusted p* = 4.17 × 10⁻⁷) and in IEZ than AEZ (log2 fold change = 16.32, *z* = 4.10, adjusted *p* = 0.00496), whereas no significant differences were detected between IEZ and BEZ **(Table S5**). The relative abundance of the dominant fungal taxa further revealed heterogeneous distributions among sampling zones **(Figure 6-A**). *Acrocalymma* showed high relative abundance in some BEZ samples but was nearly undetected in AEZ, while *Cumuliphoma* and *Vishniacozyma* showed comparatively higher abundances in AEZ. Other dominant genera, including *Pseudocercospora*, *Hannaella*, *Apiotrichum*, Trametes, and *Cutaneotrichosporon*, also exhibited variable distributions among zones. Nevertheless, when the dominant taxa were evaluated using global Kruskal–Wallis tests, none differed significantly among BEZ, IEZ, and AEZ after Benjamini–Hochberg correction (all *q* ≥ 0.450). At the family level, the abundance profiles further highlighted differences in the taxonomic composition of individual sediment samples **(Figure 6-B**). *Didymellaceae* showed the highest relative abundances in AEZ, whereas *Acrocalymmaceae* was predominantly represented in BEZ. *Bulleribasidiaceae* and *Mycosphaerellaceae* also showed comparatively higher abundances in AEZ, while *Pyrenochaetopsidaceae* and *Trichosporonaceae* were represented across multiple sampling zones.

**Figure 6.**
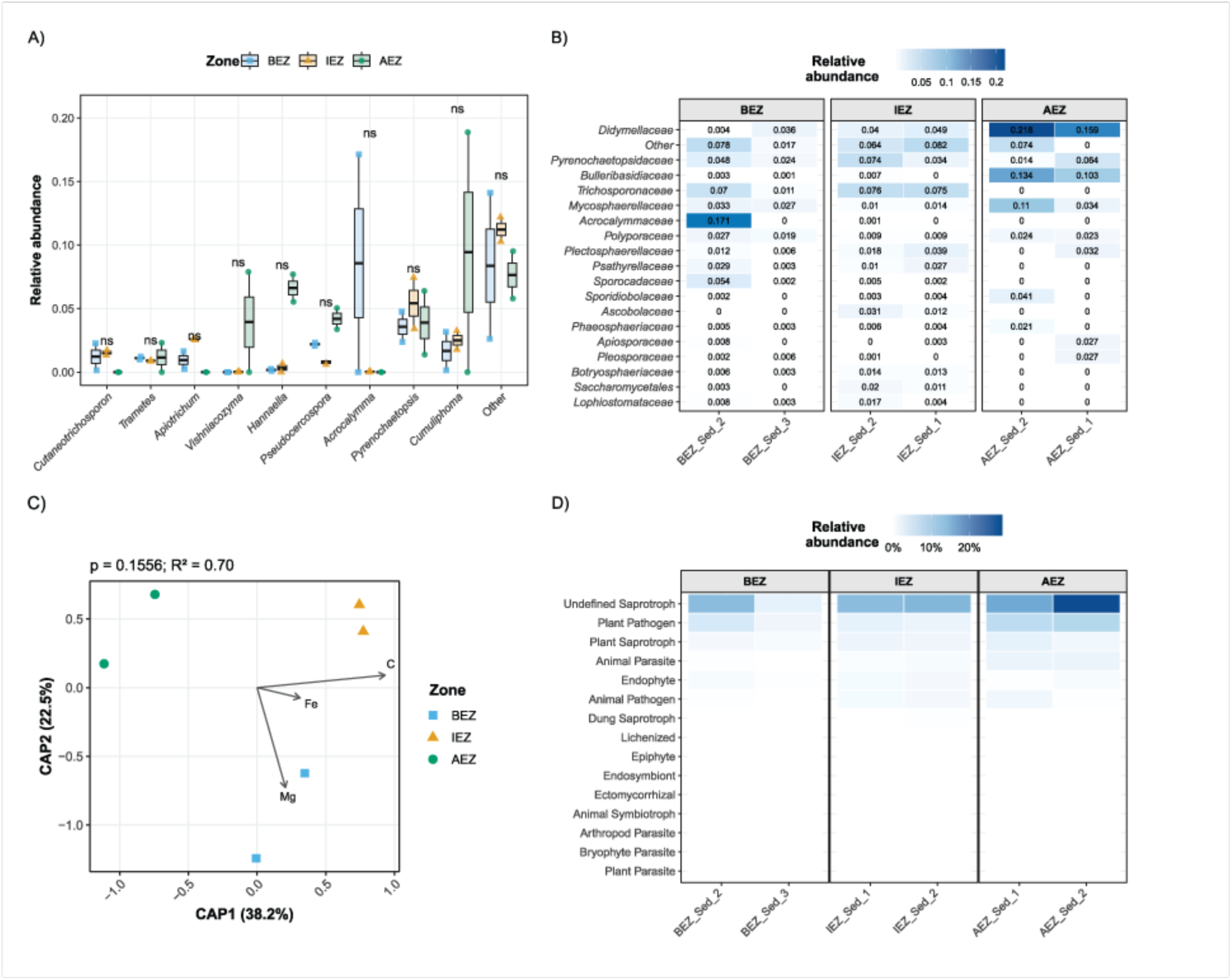
Taxonomic composition, environmental associations, and predicted ecological guilds of fungal communities in sediment. **A)** Boxplots of top genus with the highest mean relative abundance across zone (BEZ, IEZ, and AEZ), with individual observations overlaid as jittered points. Differences between zone were evaluated separately for each taxon using Wilcoxon rank-sum tests, with p-values adjusted for multiple comparisons using the Benjamini–Hochberg (BH) procedure. ns indicates q ≥ 0.05. **B**) Heatmap of the relative abundances of the most abundant fungal families across sampling zones. **C**) Distance-based redundancy analysis (dbRDA) of fungal community composition in sediment, based on Bray–Curtis dissimilarities and constrained by measured environmental variables. Points represent individual samples. Arrows show significant environmental variables (p ≤ 0.05; up to eight variables per panel), and the overall model p-value and R² are displayed above each ordination. **D**) Heatmap of predicted fungal ecological guilds in sediment samples across sampling zones. Color intensity represents the relative abundance of predicted guilds across individual samples.

### Environmental associations with fungal community

To evaluate whether variation in sediment fungal community composition was associated with the measured environmental conditions, we performed distance-based redundancy analysis (dbRDA; **Figure 6-C**). The overall environmental model was not statistically significant (*p* = 0.156), indicating that the combined set of measured variables did not significantly account for variation in fungal community composition. Among individual predictors, however, carbon was significantly associated with community structure (*p* = 0.0167), whereas magnesium and iron showed no significant associations (*p* = 0.322 and 0.475, respectively). The ordination showed a marked distribution of samples along CAP1, with AEZ samples occurring toward negative values and IEZ samples toward positive values, approximately along the carbon gradient, while BEZ samples occupied an intermediate position and were primarily differentiated along CAP2. Thus, although carbon showed an individual association with fungal community variation, the environmental model did not provide significant support for the observed community-level structuring.

### Predicted ecological guilds of fungal communities in sediment

FUNGuild-based functional annotation revealed heterogeneous distributions of predicted fungal ecological guilds across sediment samples and sampling zones **(Figure 6-D**). Undefined saprotrophs represented the most prominent functional category, particularly in AEZ, while plant pathogens and plant saprotrophs also contributed appreciably to the predicted functional profiles. Several less abundant guilds, including animal parasites, animal pathogens, endophytes, and dung saprotrophs, showed variable representation among individual samples. Despite these descriptive patterns, none of the evaluated functional categories differed significantly among BEZ, IEZ, and AEZ after Benjamini–Hochberg correction (Kruskal–Wallis tests, all adjusted *p* ≥ 0.328). Thus, the FUNGuild analysis showed variations in the relative representation of predicted ecological guilds among the sediment samples, but no statistically significant differences were observed.

## Discussion

### The geochemistry of the sediments indicates no alterations

We observed changes in different physical and chemical properties of the water samples collected from the BEZ, IAZ and AEZ. The multivariate dispersion of the data relative to the center of each group indicates a more homogeneous trend at the BEZ zone **(Figure S1)**. This response may be attributable to variables such as Ca, color, TNK, and turbidity, and could be explained by reduced disturbance in the water column. Furthermore, some variables associated with elevated nutrient levels have demonstrated a substantial increase in and/or IEZ and AEZ. The marked increase in turbidity, color, EC, TDs, and some ionic species in IEZ is consistent with patterns reported for recreational streams and waterfalls, where swimming, wading and shoreline trampling enhance sediment resuspension, particle load and organic inputs without necessarily driving classical, nutrient-saturated eutrophication (Phillip et al. 2009, Santiago et al. 2008, Lukavský et al. 2006). Similar increases in turbidity, suspended solids and conductivity associated with recreational use have been documented in streams and some tourism-impacted waters, where recreation intensity was a major predictor of water-quality degradation and ecological change (Cabello et al. 2022, Butler et al. 2021, Escarpinati et al. 2014, Turton, 2005).

In AAW, nutrient concentrations (TKN, TP) and the absence of detectable dissolved trace metals suggest that the disturbance is driven more by physical and organic loading than by chronic nutrient or metal pollution, placing this system in an early stage of water-quality alteration rather (Akhtar et al. 2021). Also, these increases suggest the addition of suspended and dissolved particles, as well as organic and minerals, possibly caused directly by anthropogenic activity in the ecotourism zone or indirectly by natural sedimentation processes that occur only in that zone (Almeida et al. 2007, Halliday et al. 2014). When comparing these results with standard regulations of Mexico (NOM-001-SEMARNAT-2021) and the United States of America (EPA, 1986), our data suggest that while parameters such as pH, alkalinity, COD, and nutrient concentrations (TNK and TP) generally comply with regulatory limits in all zones, the IEZ exceeds permissible levels of color, turbidity, and total suspended solids (TSS). Together, these results suggest that, despite detectable changes in alkalinity, the system maintains generally low levels of nutrient enrichment and metal contamination.

The elemental composition of sediments showed a consistent predominance of oxygen (45– 55%), carbon (15–20%), silicon (12–20%), calcium (15–22%), and magnesium (4–8%) across BEZ, IEZ and AEZ (**Figure 3**). These proportions are characteristic of carbonate-rich tropical fluvial sediments, formed primarily by calcium carbonate (CaCO_3_) precipitation and silicate mineral fractions, consistent with the known geology of the AAW site (SEMARNAT 2017). The predominance of Ca, Si, C, and O could indicate a sediment matrix composed of calcite/aragonite, magnesium hydroxide, and siliceous particles, a composition typical of shallow karstic systems and high-energy streams with active precipitation of CaCO_3_. The relatively low abundance of metals such as Fe, Al, Na and K shows that the system is chemically oligotrophic and not subjected to chronic metal pollution. This stability contrasts with many anthropogenically impacted lakes and rivers where sediment microbial communities are tightly coupled to gradients of metals (Zn, Cu, Cr, Cd, Pb) and phosphorus, and where these contaminants emerge as dominant drivers of bacterial community structure (Zhang et al. 2016, Wu et al. 2017, Tamburini et al. 2020, Wang et al. 2020). In the AAW, the sediments do not show increases in metals or nutrients, allowing us to rule out chemical pollution as the main cause of change. This suggests that the impact of ecotourism is mainly physical, such as the disturbance of the environment and the addition of organic matter, rather than chemical inputs.

The SEM images (**Figure S2**) support this interpretation by showing pronounced morphological disruption of sediment particles in IEZ, including fragmentation, angular edges, surface roughness and heterogeneous grain size. These features are indicative of mechanical abrasion, followed by sediment resuspension and redeposition of disturbed material. In contrast, In BEZ samples particles exhibited compact surfaces, smooth crystalline edges, and well-defined crystalline structures, indicating minimal physical disturbance and stable mineral deposition. Therefore, the results of the FE-SEM analysis support the idea that the alteration is physical, rather than chemical.

While our sampling reflected baseline conditions, seasonal fluctuations, particularly during the peak summer tourist season, are likely to amplify the observed patterns, as has been documented in similar freshwater systems (URycki et al. 2020). Furthermore, the 3 cm sediment sampling depth, while consistent with standard benthic protocols (Miranda et al. 2021), inevitably excludes deeper anaerobic layers (sediment core more than 10 cm deep). This limitation restricts our ability to characterize all vertical sediment layers and their microbial processes but also points to a clear direction for future research.

### Microbial–environment relationships across matrices

In this study, the use of primers targeting archaeal and bacterial 16S rRNA genes and the fungal ITS2 region, combined with a comprehensive analysis of the datasets, improved the characterization of microbial diversity, enabled comparisons among sites, and facilitated the identification of relationships between microbial communities and environmental factors (Martinez-Porchas et al. 2017, Peña-Ocaña et al. 2022).

The structure of the bacterial community reflects a possible trophic gradient and is consistent with findings from other freshwater systems subjected to organic enrichment (Qiu et al. 2020, Jiao et al. 2021, Betiku et al. 2021, Li et al. 2023), where functional redundancy contributes to the maintenance of ecosystem processes despite considerable taxonomic variation. The effect was even stronger for archaea, for which sampling zone accounted for a substantial proportion of compositional variation (PERMANOVA R² = 0.670). Together, these findings suggest that the spatial gradient primarily selected among microbial lineages rather than changing overall community diversity.

One of the most consistent patterns observed in the dataset was the disproportionate differentiation of AEZ in relation to BEZ and IEZ. In water, DESeq2 identified 151 differentially abundant features between BEZ and AEZ and 73 between IEZ and AEZ, compared with only one between IEZ and BEZ. The fact that this pattern reoccurs across bacteria and archaea indicates that AEZ is the primary endpoint of microbial differentiation within the studied spatial gradient. Despite being defined as distinct sampling zones (based on physiochemistry results), BEZ and IEZ generally showed substantially less taxonomic differentiation from one another. For example, Mg was more strongly associated with bacterial community than with archaeal or fungal community, a pattern that has also been reported for soil microbial communities (Yang et al. 2023).

Although microbial communities in water-impacted river systems can exhibit rapid functional responses (Bai et al. 2018, Adedire et al. 2022), our results indicate a compositional turnover associated with environmental variation. Carbon in sediment was associated with changes in both bacterial and fungal community structure, suggesting its importance as an ecological factor influencing microbial assemblages. This suggests that carbon availability is an important environmental factor influencing the composition of the microbial community in shallow oligotrophic subsurfaces (Guo et al. 2019, Bartelme et al. 2020, Da Costa et al. 2023).

The identity of several differentially abundant taxa provides additional clues regarding the ecological processes that may contribute to this differentiation. In the context of sediment bacteria, genera including *Thiobacillus*, *Flavihumibacter*, *Terrimonas*, *Sphingomonas*, and *Lacibacter* have been shown to differentiate AEZ from both BEZ and IEZ. These bacterial genera are predominantly and strictly aerobic (Justin and Kelly, 1978, Busse et al. 2003, Sheu et al. 2010, Han et al. 2017, Shameem and Sabumon 2026). In contrast, the predominant archaeal taxa, including *Methanothrix*, *Methanospirillum*, *Methanoregula*, *Nitrosarchaeum*, and *Candidatus Nitrosotenuis*, consist of anaerobic microbes that could play important roles in carbon and nitrogen transformations (Ferry and Wolfe, 1977, Oren, 2014, Zhou et al. 2023). These data suggest that oxygen availability, both acetoclastic and hydrogenotrophic methanogenesis, and nitrogen metabolism may be important determinants of ecosystem functioning; however, direct *in situ* measurements of metabolic activity are required to validate these hypotheses (Heylen et al. 2006, Urbanová et al. 2014).

The archaeal community provided a particularly informative contrast. Although taxa within the phylum Thermoplasmatota have been associated with organic pollution and methanogenic activity in surface sediments (Yang et al. 2020, Glissman et al. 2024), and archaeal communities exhibited the strongest spatial differentiation observed in this study, neither the overall dbRDA model nor any measured sediment variable explained this variation. Therefore, archaeal communities may respond to environmental gradients not captured by the measured variables. The occurrence of methanogenic and ammonia-oxidizing archaeal lineages highlights the importance of other environmental factors such as redox conditions and availability of carbon, oxygen, nitrogen-rich compounds, hydrogen and sulfur (Stahl and De La Torre 2012, Buan 2018, Baker et al. 2020). The presence of these archaeal lineages may have value as bioindicators of environmental conditions due to their specific metabolic requirements.

The detection of specialized microbial groups spanning all three domains of life, including methanogenic and ammonia-oxidizing archaea, sulfur-oxidizing bacteria, and fungi associated with organic matter degradation, highlights the complexity of microbial interactions and the potential coupling of carbon, nitrogen, sulfur, and methane cycling processes in this freshwater ecosystem. However, these findings should be interpreted with caution. PICRUSt infers potential functions from taxonomic composition but does not directly measure gene content, gene expression, or metabolic activity. Consequently, the absence of significant differences in predicted functional profiles should not be interpreted as evidence that microbial communities perform equivalent functions or that key ecosystem processes are necessarily maintained across environmental conditions. Rather, the observed taxonomic patterns provide preliminary insights into potential functional differences among zones. Conclusive assessment of functional variation and ecosystem processes within these microbial communities will require direct approaches, including multi-omics analyses and *in situ* measurements of microbial activity and biogeochemical rates.

## Conclusions and further directions

This study provides the first integrated assessment of bacterial, archaeal, and fungal communities, alongside physicochemical parameters, within the aquatic and sedimentary habitats of the freshwater stream of Agua Azul waterfalls. Our study establishes a baseline for evaluating future ecological changes in this ecosystem. Most water and sediment quality parameters remained within expected ranges; microbial diversity was largely comparable across different zones; and predicted functional profiles for bacteria, archaea, and fungi suggest that key metabolic functions are preserved.

Our findings indicate that, to date, there have been no substantial changes in microbial diversity or functional potential throughout this ecosystem. The current situation presents an opportunity to prevent eutrophication and other forms of ecosystem degradation before irreversible ecological impacts occur. In addition, we propose that Agua Azul Waterfalls could serve as a unique natural laboratory for microbial ecology, offering an opportunity to investigate microbial diversity, community assembly, and ecosystem functioning within a largely unexplored freshwater system. Further spatial and temporal studies could reveal new taxa, ecological interactions, and biogeochemical processes, which would advance our understanding of freshwater microbiomes and, in turn, guide the conservation of this type of ecosystems.

## Acknowledgments

This work was supported by StartUp funds provided to MGS-M by the Department of Molecular and Cell Biology (MCB) and the College of Liberal Arts and Sciences (CLAS) at the University of Connecticut (UConn). The authors gratefully acknowledge the additional financial support provided through the “Apoyos Únicos otorgados a los integrantes del Sistema Estatal de Investigadores 2023” program, funded by the Instituto de Ciencia, Tecnología e Innovación del Estado de Chiapas (ICTIECH), now the Agencia Digital Tecnológica. The support of our institutions was fundamental to the execution of this study and the contributions to the Agua Azul community.

The authors thank Alexander Neil Poulter, a member of the UConn Microbial Ecophysiology Laboratory, for his help in preparing the samples for sequencing, and Dr. Kendra Maas (UConn CGI) for her assistance in primer design and sample processing.

The authors thank the community of Agua Azul (Chiapas, Mexico) for their support and guidance. Their cooperation was essential to the development of this project. We additionally acknowledge Jesús Antonio Domínguez-Espinosa for his valuable assistance in coordinating and securing the permissions with the local community.

To the local Indigenous community (text in tseltal): Te mach’atic la spasic ja’ iproyecto in to, ya calbetic bayal wocol te lum Agua azul sbiil (Chiapas, Mexico) yu’un te muc’ul coltayel la yac’botic, soc ja’ nix me jich yu’un te bit’il la yac’botic ts’ajc’tic te slum-sq’uinalic ta swenta yu’un te muestra. Jich yu’un mu’c scloplal ja’ c’ot c’alal la xjajch’ ta spasel ja’ i proyecto in to.

We, the authors, belong to Indigenous communities, Zoque (*O’depüt*), Olmeca (*Olmecatl*), Zapoteca (*Binnizá*) and Mixteca (*Ñuu Savi*), and we thank our people and ancestors for their strength and resilience in protecting the land, traditions, and knowledge.

## Funding

This work did not receive external funding.

## Data and code availability

Raw sequence data and associated metadata have been deposited in the NCBI Sequence Read Archive (SRA) under BioProject accession PRJNA1425486, including bacterial and archaeal 16S rRNA gene amplicons and fungal ITS sequences. The complete bioinformatic workflow, parameter files, and downstream R analysis scripts are publicly available at https://github.com/OvADN2347/ecotourism-microbiota.

## Statement on the use of artificial intelligence (AI) tools

ChatGPT (OpenAI) was used to assist with coding and data-querying tasks. All code, analyses, and outputs generated with AI assistance were reviewed and validated by the authors. The authors take full responsibility for the content of this manuscript.

## Supplemental material

Supplementary materials are included (Figure S1-S6 and Supplementary Table 1-6). Additional information regarding the sequencing and processing of the samples can be found in the repository.

## GLOSSARY

AAW: Agua Azul Waterfall
AEZ: After Ecotourism Zone
ANOVA: Analysis of Variance
APHA: American Public Health Association
ASV: Amplicon Sequence Variants
BEZ: Before Ecotourism Zone
CaCO_3_: Calcium Carbonate
COD: Chemical Oxygen Demand
CONANP: National Commission of Natural Protected Areas (Mexico) Comisión Nacional de Áreas Naturales Protegidas (México)
DNA: Deoxyribonucleic Acid
EC: Electrical Conductivity
EDS: Energy-Dispersive X-ray Spectroscopy
EPA: Environmental Protection Agency
FE-SEM: Field-Emission Scanning Electron Microscope
ICP-OES: Inductively Coupled Plasma Optical Emission Spectrometry
IEZ: In Ecotourism Zone
ITS: Internal Transcribed Spacer
OTU: Operational Taxonomic Unit
Pt-Co: Platinum-Cobalt color scale
PERMOVA: Permutational Multivariate Analysis of Variance
SEMARNAT: Ministry of Environment and Natural Resources (Mexico) Secretaría del Medio Ambiente y Recursos Naturales (México)
TDS: Total Dissolved Solids
TKN: Total Kjeldahl Nitrogen
TP: Total Phosphorus
TS: Total Solids
TSS: Total Suspended Solids
TVS: Total Volatile Solids

## Supplementary Material

**Figure S1.** Principal component analysis of water physicochemical characteristics across sampling zones.

**Figure S2.** Surface morphology of sediment samples from the Agua Azul Waterfalls obtained by field-emission scanning electron microscopy (FE-SEM).

**Figure S3.** Microbial community diversity and taxonomic composition for bacteria in water samples.

**Figure S4.** Microbial community diversity and taxonomic composition for bacteria in sediment.

**Figure S5.** Microbial community diversity and taxonomic composition for archaea in sediment samples.

**Figure S6.** Microbial community diversity and taxonomic composition for fungal community in sediments samples.

***Note:*** The following files will be available once the manuscript is published in a peer-reviewed journal. If you need access to the raw data before that date, please contact Dr. Geo Santiago-Martínez.

**Supplementary Table 1:** Raw data of physicochemical determinations in water and sediment.

**Supplementary Table 2:** Differential abundance analysis of bacterial genera across sampling zones in water samples.

**Supplementary Table 3:** Differential abundance analysis of bacterial genera across sampling zones in sediment samples.

**Supplementary Table 4:** Differential abundance analysis of archaeal genera across sampling zones in sediment samples.

**Supplementary Table 5:** Differential abundance analysis of fungal genera across sampling zones in sediment samples.

## Notes

### Competing Interest Statement

The authors have declared no competing interest.

### Summary of Updates

Some references were added and others were corrected. The length of the manuscript was shortened. The quality of the figures was improved. The figures and tables from the supplementary material were included in the main text.

## References

1. Abarenkov K, Zirk A, Piirmann T et al. UNITE QIIME release for Fungi. Version 04.04.2024. UNITE Community, 2024.

2. Adedire DE, Jimoh AO, Kashim-Bello Z et al. Microbiome diversity analysis of the bacterial community in Idah River, Kogi State, Nigeria. Adv Microbiol 2022;12:343–62.

3. Akhtar N, Syakir Ishak MI, Bhawani SA, Umar K. Various natural and anthropogenic factors responsible for water quality degradation: a review. Water 2021;13:2660.

4. Almeida CA, Quintar S, González P, Mallea MA. Influence of urbanization and tourist activities on the water quality of the Potrero de los Funes River (San Luis, Argentina). Environ Monit Assess 2007;133:459–65.

5. American Public Health Association, (APHA). American Water Works Association, Water Environment Federation. Standard Methods for the Examination of Water and Wastewater. 22nd edn. Washington, DC: American Public Health Association, 2012.

6. Anderson MJ, Ellingsen KE, McArdle BH. Multivariate dispersion as a measure of beta diversity. Ecol Lett 2006;9:683–93.

7. Aparici-Carratalá D, Esclapez J, Bautista V et al. Archaea: current and potential biotechnological applications. Res Microbiol 2023;174:104080.

8. Bai M, Dong F, Jia Y et al. Impact of sediment plume on benthic microbial community in deep-sea mining. Water 2025;17:3013.

9. Bai Y, Wang Q, Liao K et al. Fungal community as a bioindicator to reflect anthropogenic activities in a river ecosystem. Front Microbiol 2018;9:315.

10. Baker BJ, De Anda V, Seitz KW, Dombrowski N, Santoro AE, Lloyd KG. Diversity, ecology and evolution of Archaea. Nat Microbiol 2020;5:887–900.

11. Bartelme RP, Custer JM, Dupont CL, Espinoza JL, Torralba M, Khalili B, Carini P. Influence of substrate concentration on the culturability of heterotrophic soil microbes isolated by high-throughput dilution-to-extinction cultivation. mSphere 2020;5:e00624–19.

12. Betiku OC, Sarjeant KC, Ngatia LW et al. Evaluation of microbial diversity of three recreational water bodies using 16S rRNA metagenomic approach. Sci Total Environ 2021;771:144773.

13. Buan NR. Methanogens: pushing the boundaries of biology. Emerg Top Life Sci 2018;2:629–46.

14. Busse HJ, Denner EBM, Buczolits S et al. *Sphingomonas aurantiaca* sp. nov., *Sphingomonas aerolata* sp. nov. and *Sphingomonas faeni* sp. nov., air-and dustborne and Antarctic, orange-pigmented, psychrotolerant bacteria, and emended description of the genus *Sphingomonas*. Int J Syst Evol Microbiol 2003;53:1253–60.

15. Butler B, Pearson RG, Birtles RA. Water-quality and ecosystem impacts of recreation in streams: monitoring and management. Environ Chall 2021;5:100328.

16. Cabello CA, Canini ND, Lluisma BC. Water quality assessment of Dodiongan Falls in Bonbonon, Iligan City, Philippines. AIMS Environ Sci 2022;9:526–37.

17. Comisión Nacional de Áreas Naturales Protegidas (CONANP). Ficha del Sistema de Información, Monitoreo y Evaluación para la Conservación (SIMEC) de Cascadas de Agua Azul. Gobierno de México, 2025. https://simec.conanp.gob.mx/ficha.php?anp=130&reg=11 (28 November 2026, date last accessed).

18. da Costa C, Colin Y, Debret M et al. Shifts in sediment bacterial communities reflect changes in depositional environments in a fluviatile context. Sci Total Environ 2023;885:163890.

19. Douglas GM, Maffei VJ, Zaneveld JR et al. PICRUSt2 for prediction of metagenome functions. Nat Biotechnol 2020;38:685–88.

20. Escarpinati SC, Siqueira T, Medina PB, Roque FO. Short-term effects of visitor trampling on macroinvertebrates in karst streams in an ecotourism region. Environ Monit Assess 2014;186:1655–63.

21. Ferry JG, Wolfe RS. Nutritional and biochemical characterization of *Methanospirillum hungatii*. Appl Environ Microbiol 1977;34:371–76.

22. Garza-Tovar JR, Sánchez-Crispín A, Figueroa-Encino A. Territorial structure of tourism in the region Palenque-Cascadas de Agua Azul, Chiapas, Mexico. Dimens Tur 2020;4:63–90.

23. Glissman K, Chin KJ, Casper P, Conrad R. Methanogenic pathway and archaeal community structure in the sediment of eutrophic Lake Dagow: effect of temperature. Microb Ecol 2004;48:389–99.

24. Guo XP, Yang Y, Niu ZS et al. Characteristics of microbial community indicate anthropogenic impact on the sediments along the Yangtze Estuary and its coastal area, China. Sci Total Environ 2019;648:306–14.

25. Halliday E, McLellan SL, Amaral-Zettler LA et al. Comparison of bacterial communities in sands and water at beaches with bacterial water quality violations. PLoS One 2014;9:e90815.

26. Han JH, Baek K, Lee MH. *Lacibacter nakdongensis* sp. nov., isolated from river sediment. Int J Syst Evol Microbiol 2017;67:352–56.

27. Heylen K, Gevers D, Vanparys B et al. The incidence of *nirS* and *nirK* and their genetic heterogeneity in cultivated denitrifiers. Environ Microbiol 2006;8:2012–21.

28. Ihrmark K, Bödeker ITM, Cruz-Martinez K et al. New primers to amplify the fungal ITS2 region: evaluation by 454-sequencing of artificial and natural communities. FEMS Microbiol Ecol 2012;82:666–77.

29. Jiao S, Peng Z, Qi J et al. Linking bacterial-fungal relationships to microbial diversity and soil nutrient cycling. mSystems 2021;6:e00112-21.

30. Justin P, Kelly DP. Growth kinetics of *Thiobacillus denitrificans* in anaerobic and aerobic chemostat culture. J Gen Microbiol 1978;107:123–30.

31. Kozich JJ, Westcott SL, Baxter NT et al. Development of a dual-index sequencing strategy and curation pipeline for analyzing amplicon sequence data on the MiSeq Illumina sequencing platform. Appl Environ Microbiol 2013;79:5112–20.

32. Kyle KE, Klassen JL. Untrimmed ITS2 metabarcode sequences cause artificially reduced abundances of specific fungal taxa. Appl Environ Microbiol 2025;91:e01537–24

33. Li X, Meng Z, Chen K et al. Comparing diversity patterns and processes of microbial community assembly in water column and sediment in Lake Wuchang, China. PeerJ 2023;11:e14592.

34. Libert-Amico A. Scenic beauty and territorial conflicts: manufacturing nature in the Agua Azul Waterfalls, Chiapas. EntreDiversidades 2019;6:9–42.

35. Lukavský J, Moravcová A, Nedbalová L, Rauch O. Phytobenthos and water quality of mountain streams in the Bohemian Forest under the influence of recreational activity. Biologia 2006;61(Suppl 20):S533–S542.

36. Martinez-Porchas M, Villalpando-Canchola E, Suarez LEO, Vargas-Albores F. How conserved are the conserved 16S rRNA regions? PeerJ 2017;5:e3036.

37. McAllister TG, Walker LA, Smiles ND, Jennings L, Moggridge B, Naepi S, et al. Decolonize scientific institutions, don’t just diversify them. Nature 2025;644:331–34.

38. Miranda LS, Ayoko GA, Egodawatta P et al. Physico-chemical properties of sediments governing the bioavailability of heavy metals in urban waterways. Sci Total Environ 2021;763:142984.

39. Nguyen NH, Song Z, Bates ST et al. FUNGuild: an open annotation tool for parsing fungal community datasets by ecological guild. Fungal Ecol 2016;20:241–48.

40. Oren A. The family Methanoregulaceae. In: Rosenberg E, DeLong EF, Lory S, Stackebrandt E, Thompson F, eds. The Prokaryotes: Other Major Lineages of Bacteria and the Archaea. Berlin, Heidelberg: Springer, 2014, 253–58.

41. Peña-Ocaña BA, Ovando-Ovando CI, Puente-Sánchez F et al. Metagenomic and metabolic analyses of poly-extreme microbiome from an active crater volcano lake. Environ Res 2022;203:111862.

42. Phillip DAT, Antoine P, Cooper V et al. Impact of recreation on recreational water quality of a small tropical stream. J Environ Monit 2009;11:1192–98.

43. Qiu H, Gu L, Sun B et al. Metagenomic analysis revealed that terrestrial pollutants override the effects of seasonal variation on microbiome in river sediments. Bull Environ Contam Toxicol 2020;105:892–98.

44. Ramírez-Galdámez RC, Villalobos-Maldonado JJ, Cruz-Salomón A et al. Efficient bioreactor system for biological treatment and methane production from fish processing wastewater. J Water Process Eng 2024;68:106476.

45. Santiago LE, Gonzalez-Caban A, Loomis J. A model for predicting daily peak visitation and implications for recreation management and water quality: evidence from two rivers in Puerto Rico. Environ Manage 2008;41:904–14.

46. Secretaría de Medio Ambiente y Recursos Naturales (SEMARNAT). Cascada de Agua Azul, un paraíso protegido por el gobierno de la República. Mexico City: SEMARNAT, 2017. https://www.gob.mx/semarnat/articulos/cascada-de-agua-azul-un-paraiso-protegido-por-elgobierno-de-la-republica (28 November 2026, date last accessed).

47. Secretaría de Medio Ambiente y Recursos Naturales (SEMARNAT). NOM-059-SEMARNAT-2010: Environmental Protection, Mexican Official Standard, Protection of native species of flora and fauna in Mexico, risk categories and specifications for inclusion, exclusion or change, list of species at risk. Mexico City: SEMARNAT, 2010.

48. Secretaría de Medio Ambiente y Recursos Naturales (SEMARNAT). NOM-001-SEMARNAT-2021: Environmental Protection, Mexican Official Standard, permissible limits of pollutants in wastewater discharges to receiving waters owned by the nation. Mexico City: SEMARNAT, 2021.

49. Shameem KS, Sabumon PC. Development of aerobic micro granular sludge in a sequencing batch airlift reactor for simultaneous removal of COD, NH₄⁺, PO₄³⁻ and SO₄²⁻. Chem Eng J Adv 2026;25:101006.

50. Sheu SY, Cho NT, Arun AB, Chen WM. *Terrimonas aquatica* sp. nov., isolated from a freshwater spring. Int J Syst Evol Microbiol 2010;60:2705–09.

51. Stahl DA, De La Torre JR. Physiology and diversity of ammonia-oxidizing archaea. Annu Rev Microbiol 2012;66:83–101.

52. Tamburini E, Doni L, Lussu R et al. Impacts of anthropogenic pollutants on benthic prokaryotic communities in Mediterranean touristic ports. Front Microbiol 2020;11:1234.

53. Turton SM. Managing environmental impacts of recreation and tourism in rainforests of the Wet Tropics of Queensland World Heritage Area. Geogr Res 2005;43:140–51.

54. Urbanová Z, Bárta J. Microbial community composition and in silico predicted metabolic potential reflect biogeochemical gradients between distinct peatland types. FEMS Microbiol Ecol 2014;90:633–46.

55. Urycki DR, Good SP, Crump BC et al. River microbiome composition reflects macroscale climatic and geomorphic differences in headwater streams. Front Water 2020;2:574728.

56. US Environmental Protection Agency (EPA). Quality Criteria for Water. Washington, DC: Office of Water Regulations and Standards, 1986.

57. US Environmental Protection Agency (EPA). Sampling and Consideration of Variability (Temporal and Spatial) for Monitoring of Recreational Waters (EPA 823-R-10-005). Washington, DC: Office of Water, 2010.

58. Wang W, Yang Y, Zhou Y et al. Impact of anthropogenic activities on the sediment microbial communities of Baiyangdian shallow lake. Int J Sediment Res 2020;35:180–92.

59. Wu W, Dong C, Wu J et al. Ecological effects of soil properties and metal concentrations on the composition and diversity of microbial communities associated with land use patterns in an electronic waste recycling region. Sci Total Environ 2017;601-602:57-65.

60. Yang W, Ji Z, Wu A et al. Inconsistent responses of soil bacterial and fungal community diversity and network structure to magnesium fertilization in tea (*Camellia sinensis*) plantation soils. Appl Soil Ecol 2023;191:105055.

61. Yang Y, Chen J, Tong T et al. Influences of eutrophication on methanogenesis pathways and methanogenic microbial community structures in freshwater lakes. Environ Pollut 2020;260:114106.

62. Zhang C, Nie S, Liang J et al. Effects of heavy metals and soil physicochemical properties on wetland soil microbial biomass and bacterial community structure. Sci Total Environ 2016;557-558:785-90.

63. Zhou J, Smith JA, Li M, Holmes DE. Methane production by *Methanothrix thermoacetophila* via direct interspecies electron transfer with *Geobacter metallireducens*. mBio 2023;14:e00360–23.

