## Supplemental FigS1-S6 for "Impact of ecotourism on the diversity of bacteria, archaea, and fungi in the freshwater stream of the Agua Azul Waterfalls in southeastern Mexico"

[1]Escuela de Ciencias Químicas, Universidad Autónoma de Chiapas (UNACH), Ocozocoautla de Espinosa 29140, Chiapas, México.
[2]Department of Molecular and Cell Biology, The University of Connecticut (UConn), Storrs 06269, Connecticut, United States.
[3]Sección de Electrónica de Estado Sólido-Ingeniería Eléctrica (SEES), CINVESTAV- IPN, San Pedro Zacatenco 07360, Ciudad de México, México.

[†] Contributed equally to this study. Author order was determined alphabetically.

**[#] Address correspondence to:**
Abumalé Cruz-Salomón, PhD.
Michel Geovanni Santiago-Martínez, PhD.

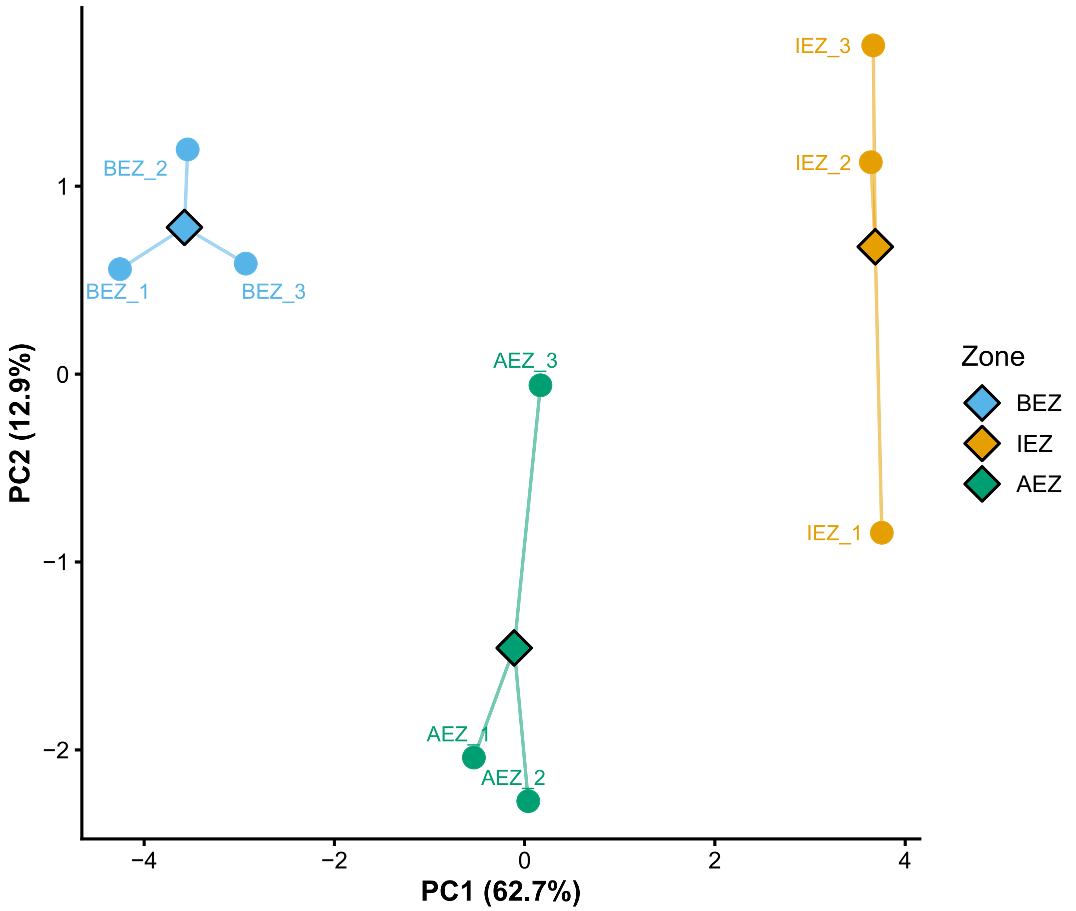


**Figure S1. Principal component analysis of water physicochemical characteristics across sampling zones.** PCA was performed on centered and standardized physicochemical variables to visualize multivariate variation among water samples across zone (BEZ, IEZ, and AEZ). Points represent individual samples (n = 3), and black-outlined diamonds represent the centroid of each sampling zone in the PC1–PC2 ordination space.

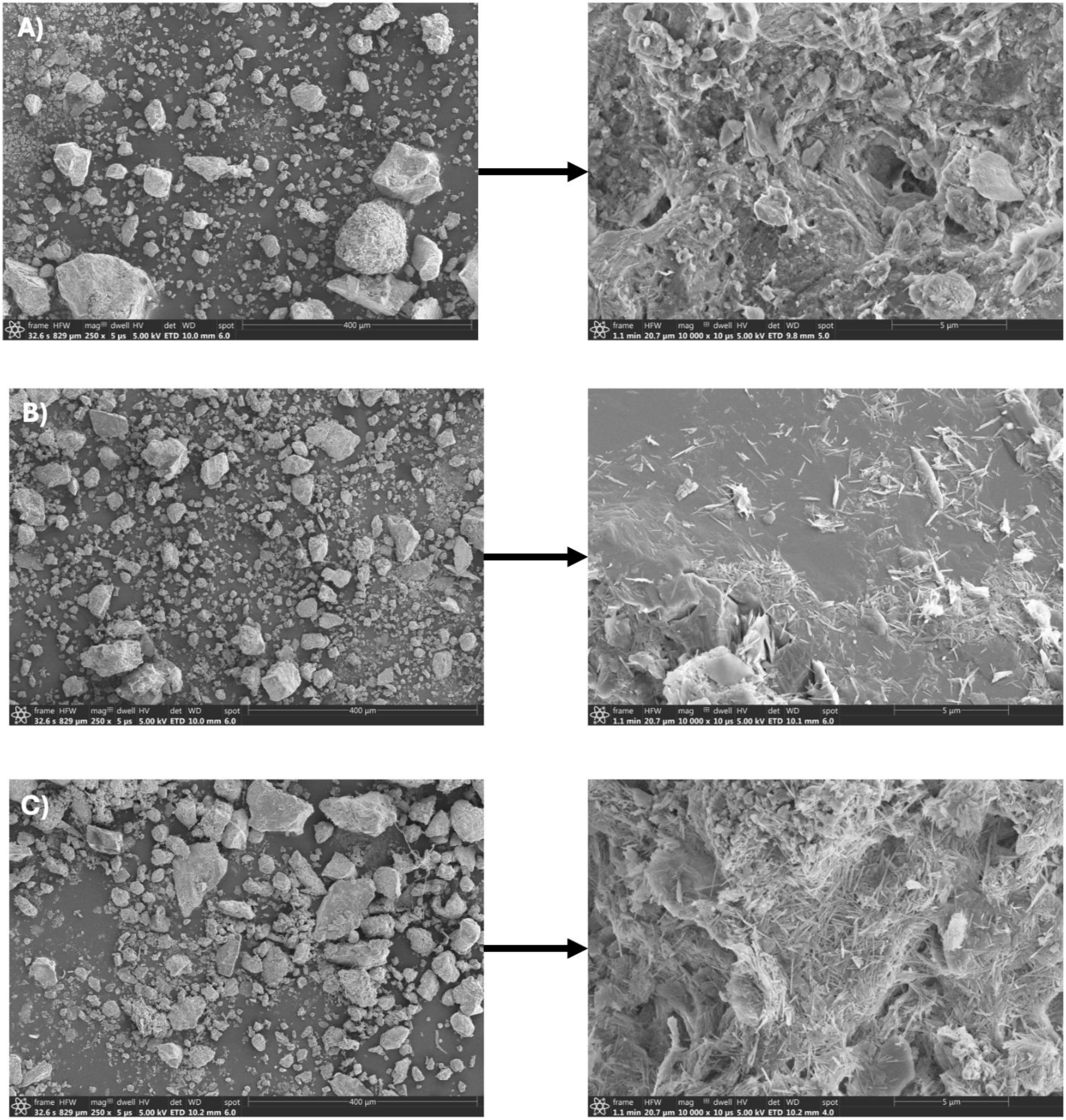


**Figure S2. Surface morphology of sediment samples from the Agua Azul Waterfalls obtained by field-emission scanning electron microscopy (FE-SEM).** (a) before the ecotourism zone (BEZ), (b) in the ecotourism zone (IEZ), and (c) after the ecotourism zone (AEZ). Magnifications: left panels ≈ 250× (scale ≈ 400 μm); right panels ≈ 10,000× (scale ≈ 5 μm).

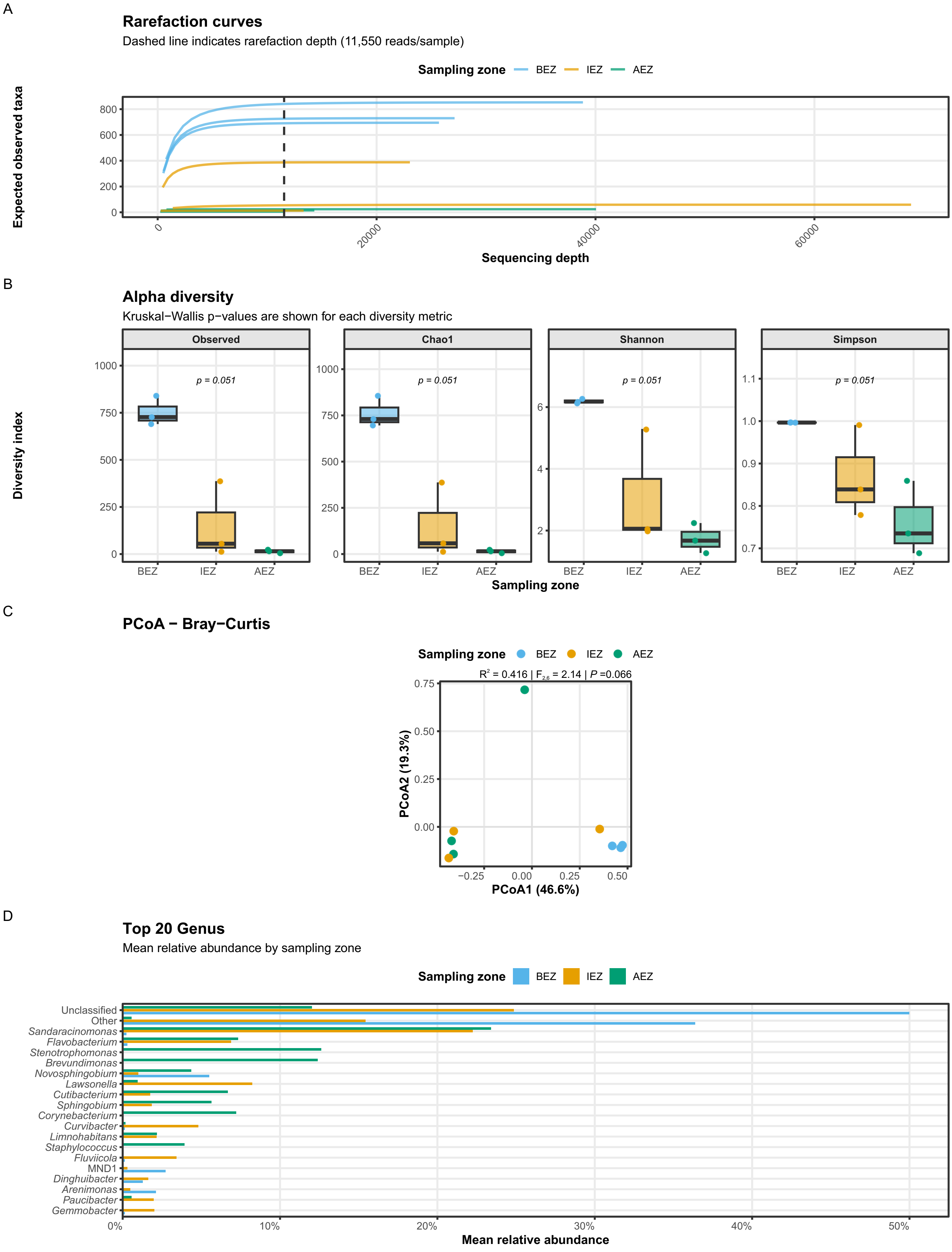


**Figure S3. Microbial community diversity and taxonomic composition for bacteria in water samples. A)** Sample-based rarefaction curves showing expected observed taxonomic richness as a function of sequencing depth. Curves shows the zone and the dashed vertical line indicates the rarefaction depth (11,550 reads per sample). **B)** Alpha-diversity estimates across sampling groups. Points represent individual samples and boxes summarize the distribution within each group. **C)** Principal coordinates analysis (PCoA) based on Bray-Curtis dissimilarities. **D)** Mean relative abundance of the most abundant genera across sampling groups. Genera selected and ranked according to their overall mean relative abundance, and bars show the mean relative abundance within each sampling group.

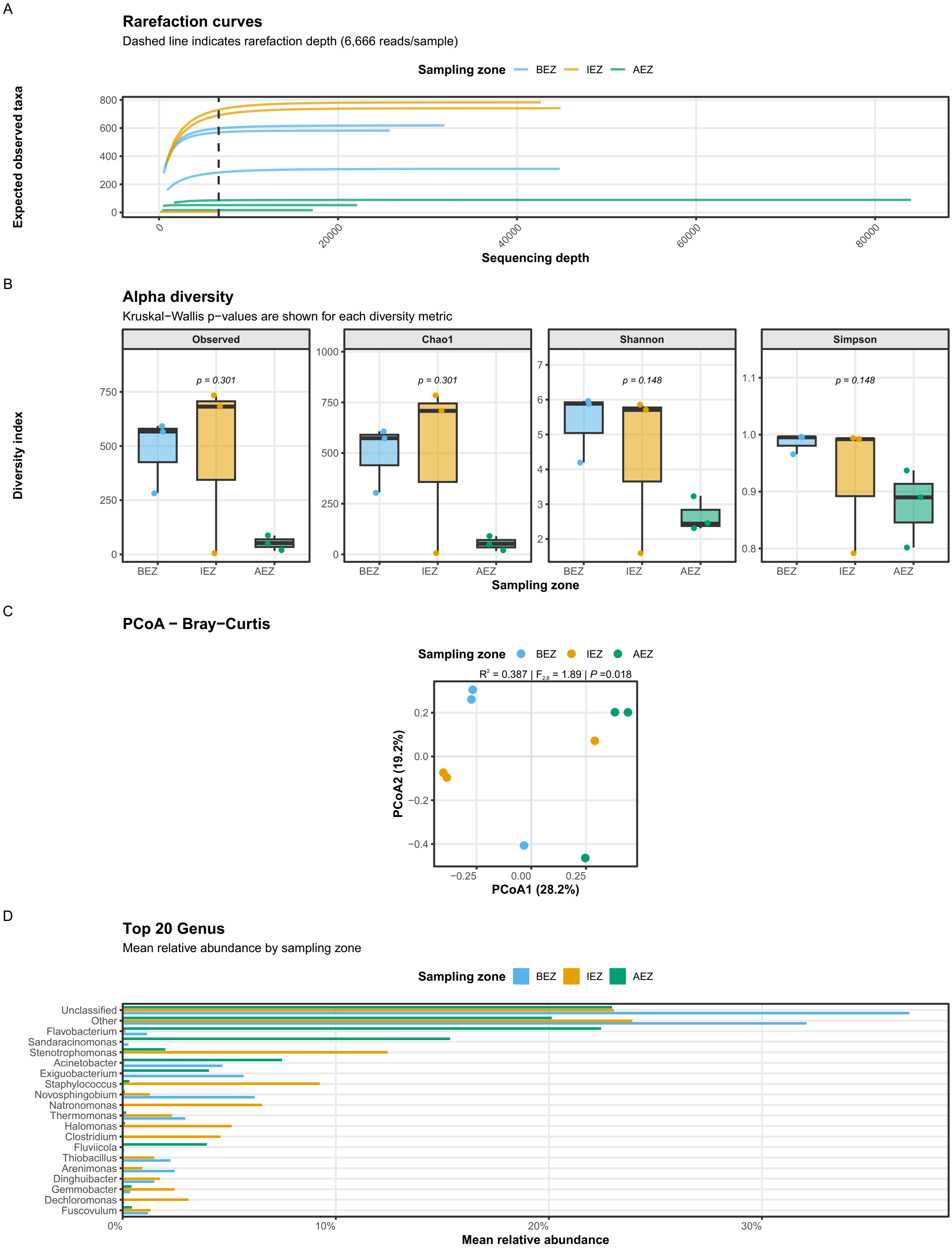


**Figure S4. Microbial community diversity and taxonomic composition for bacteria in sediment. A)** Sample-based rarefaction curves showing expected observed taxonomic richness as a function of sequencing depth. Curves show the zone and the dashed vertical line indicates the rarefaction depth (6,666 reads per sample). **B)** Alpha-diversity estimates across sampling groups. Points represent individual samples and boxes summarize the distribution within each group. **C)** PCoA based on Bray-Curtis dissimilarities. **D)** Mean relative abundance of the most abundant genera across sampling groups. Genera selected and ranked according to their overall mean relative abundance Bars show the mean relative abundance within each sampling group.

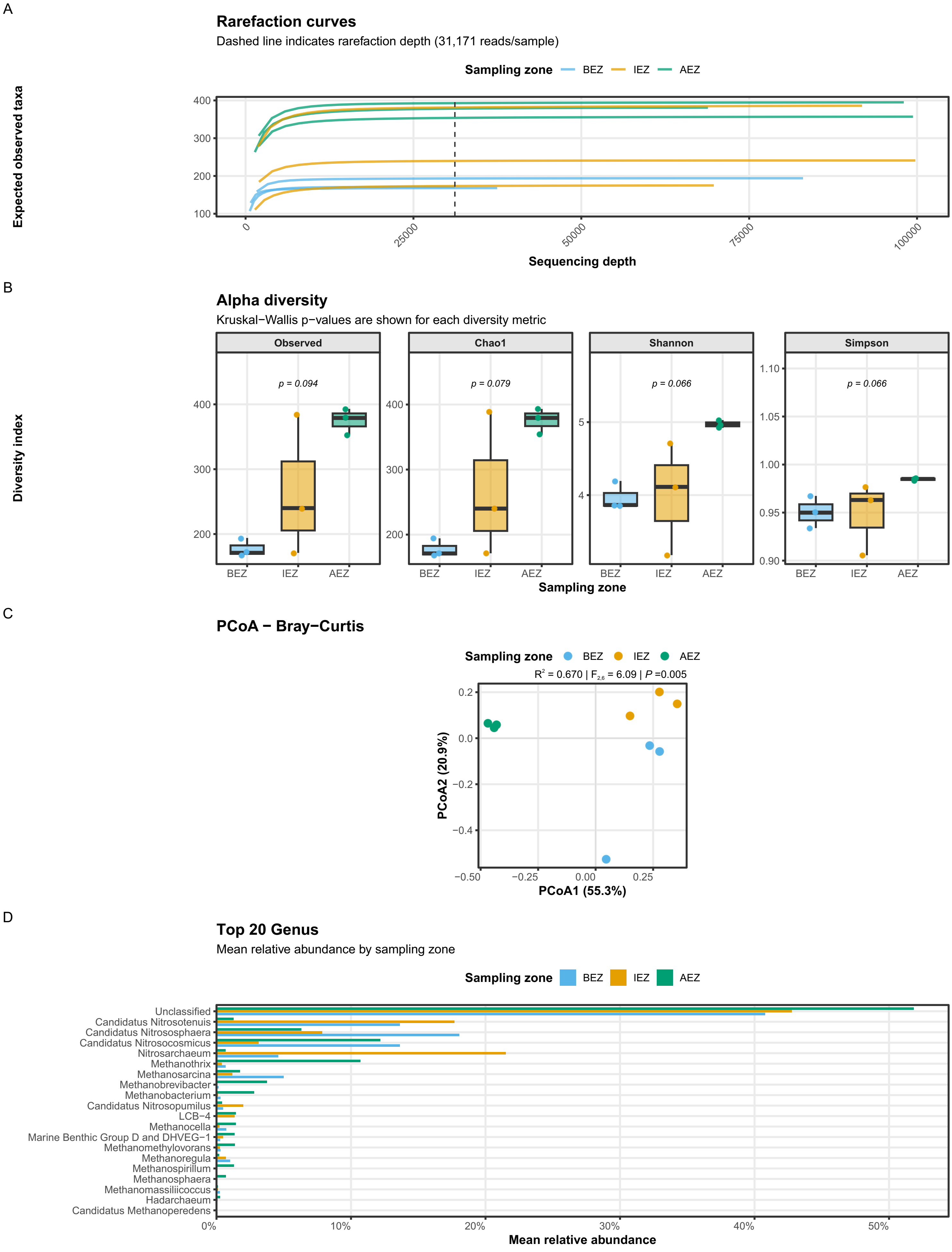


**Figure S5. Microbial community diversity and taxonomic composition for archaea in sediment samples. A)** Sample-based rarefaction curves showing expected observed taxonomic richness as a function of sequencing depth. Dashed vertical line indicates the rarefaction depth used for downstream diversity analyses (31,171 reads per sample). **B)** Alpha-diversity estimates across sampling groups. Points represent individual samples and boxes summarize the distribution within each group. **C)** PCoA based on Bray-Curtis dissimilarities. Points represent individual samples colored according to zone. **D)** Mean relative abundance of the most abundant genera across sampling groups. Genera selected and ranked according to their overall mean relative abundance, and bars show the mean relative abundance within each sampling group.

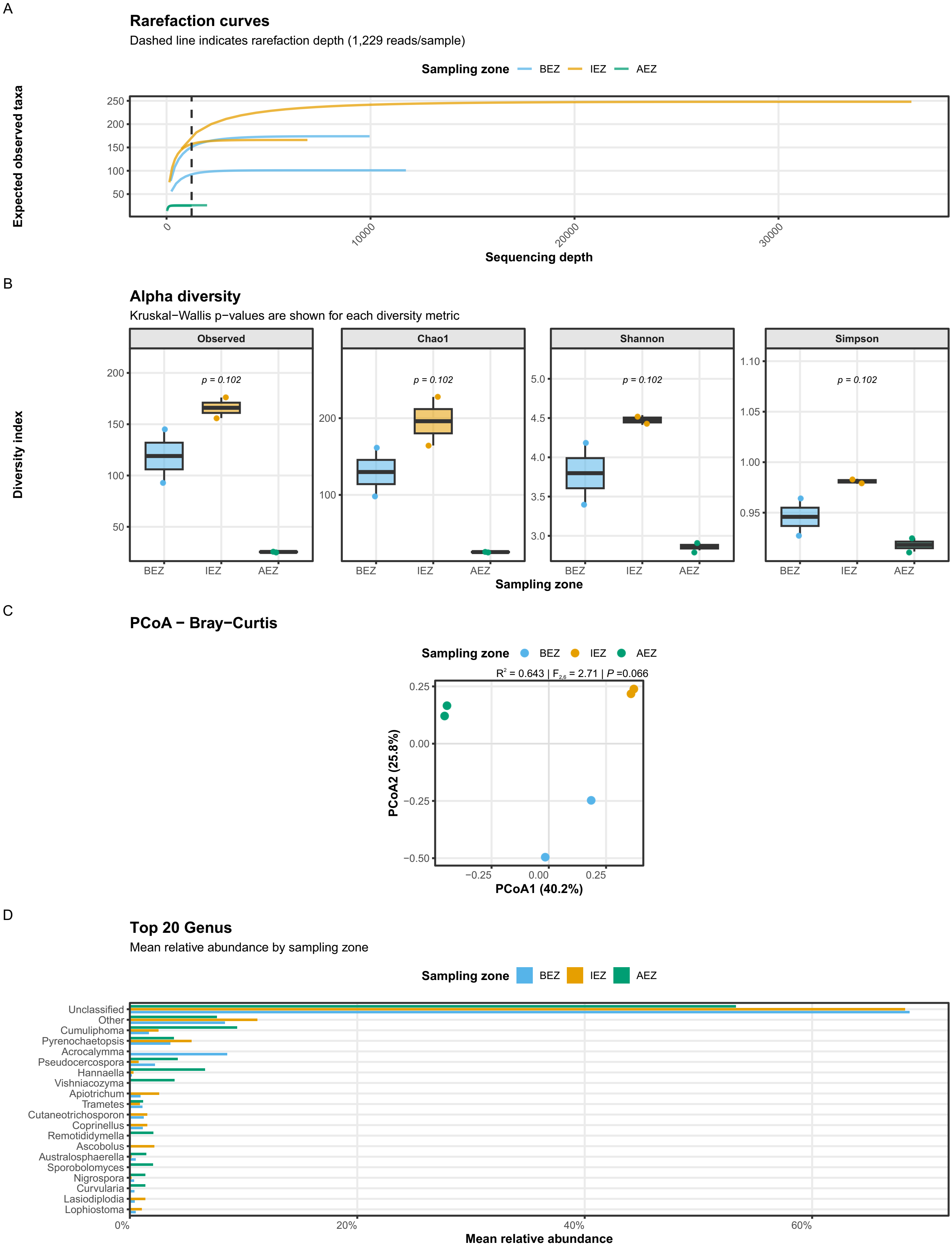


**Figure S6. Microbial community diversity and taxonomic composition for fungal community in sediments samples. A)** Sample-based rarefaction curves showing expected observed taxonomic richness. Curves show zone and dashed vertical line indicates the rarefaction depth. **B)** Alpha-diversity estimates across sampling groups. Points represent individual samples and boxes summarize the distribution within each group. **C)** PCoA based on Bray-Curtis dissimilarities. **D)** Mean relative abundance of the most abundant genera across sampling groups.
